# Remifentanil self-administration promotes circuit- and sex-specific adaptations within the prefrontal-accumbens pathways

**DOI:** 10.64898/2026.03.21.713428

**Authors:** Saurabh S. Kokane, Shalana I. Atwell, Aric C. Madayag, Eden M. Anderson, Skyler Demis, Annabel Engelhardt, Logan Friedrich, Matthew C. Hearing

## Abstract

The nucleus accumbens (NAc) and its excitatory input from the medial prefrontal cortex (mPFC) form a critical circuit underlying drug-induced plasticity associated with addiction-related behaviors. However, baseline differences in excitatory signaling across NAc subcircuits and sex-specific neuroadaptations following opioid self-administration remain poorly understood. Here, we examined synaptic signaling in mPFC–NAc pathways in drug-naïve mice and after abstinence from remifentanil self-administration. Under drug-naïve conditions, AMPA receptor– mediated glutamatergic signaling was generally elevated in D2 medium spiny neurons (MSNs) of both the NAc core and shell across sexes, while females exhibited greater excitatory signaling in D1 MSNs of the NAc core compared with males. Pathway-specific analyses revealed that prelimbic cortex (PL) inputs to NAc core D2 MSNs displayed enhanced calcium-permeable AMPA receptor (CP-AMPAR) signaling and increased presynaptic release relative to D1 MSNs. Following abstinence from remifentanil self-administration, miniature excitatory postsynaptic current analyses showed increased excitatory drive at D1 MSNs and decreased drive at D2 MSNs, largely restricted to the NAc core. At PL–Core D1 MSN synapses, remifentanil reduced AMPA/NMDA ratios, consistent with increased CP-AMPAR incorporation in males and females, while increasing presynaptic signaling exclusively in males. In contrast, PL-Core D2 MSN synapses showed a reduction in presynaptic signaling across sex, while ostensibly weakening postsynaptic signaling selectively in males through reductions in CP-AMPAR signaling. At infralimbic cortex (IL)–shell inputs, a reduction in AMPAR rectification indices at D1 MSN synapses was produced by remifentanil, while release probability was decreased at D2 MSN synapses in males only. Together, these findings reveal sex- and pathway-specific synaptic adaptations within mPFC–NAc circuits that may be obscured by global measures of excitatory transmission and identify baseline circuit differences that may shape opioid-induced plasticity.

## 1 INTRODUCTION

Opioid use disorder (OUD) develops from deficits in cognitive control that escalate opioid intake, strengthen drug-seeking behaviors, and increase an individual’s propensity to relapse. Long-lasting molecular and physiological neuroadaptations in key nodes of the reward pathway form the molecular and cellular basis of these maladaptive behaviors (Koob *et al*., 2004; Koob & Volkow, 2016). Clinical findings indicate that dysfunction of the prefrontal cortex (PFC) emerges across substance use disorders (Koob & Volkow, 2010). The rodent medial PFC (mPFC), a region homologous to the primate PFC, plays a key role in governing numerous aspects of drug reward- and drug seeking behaviors (Everitt & Robbins, 2005; Koob & Volkow, 2010). The mPFC receives and sends input to multiple limbic and midbrain targets implicated in opioid relapse. Importantly, extensive anatomical and causal evidence demonstrates that divergent mPFC-NAc projections exert robust, bidirectional control over opioid seeking (LaLumiere & Kalivas, 2008; Rogers *et al*., 2008; Van den Oever *et al*., 2010; Shen *et al*., 2011; Bossert *et al*., 2012; Marchant *et al*., 2016; Hearing, 2019). Given the role of the mPFC and its interconnected nature, understanding how this region and its subcircuits are altered by opioids will provide a mechanistically precise and translationally relevant framework for understanding cortical control of opioid seeking.

The mPFC-NAc pathway can be divided into two primary projections consisting of the prelimbic cortex projection to the core region of the NAc (PL-Core) and infralimbic cortical projections to the shell subdivision (IL-Shell) (Sesack *et al*., 1989; Berendse, 1992; Brog *et al*., 1993; Heidbreder & Groenewegen, 2003; Heimer, 2003; Vertes, 2004; Hoover & Vertes, 2007). Numerous lines of research to date have demonstrated the necessity of the PL to NAc core (PL-Core) pathway in many different types of reinstatement to drug-seeking (McFarland *et al*., 2003; LaLumiere & Kalivas, 2008; Rogers *et al*., 2008). Alternatively, while the IL-Shell pathway has been noted to inhibit psychostimulant seeking, evidence supports a potential role in promoting heroin seeking (Rogers *et al*., 2008; Van den Oever *et al*., 2010; Bossert *et al*., 2012; Peters *et al*., 2013; Hearing *et al*., 2016; Marchant *et al*., 2016). . Recent findings have identified opioid-induced adaptations in mPFC output neurons (Anderson *et al*., 2021), including those projecting to the NAc (Kokane *et al*., 2023) that are causally linked to impairments in cognition and opioid-seeking, respectively. However, what remains unknown is how these adaptations translate to plasticity downstream in the NAc to change behavior.

Like subregions of the medial PFC, the NAc core and shell show divergent responses to non-contingent morphine with dissociable and overlapping roles in opioid reward and seeking behavior (Zahm & Brog, 1992; Pontieri *et al*., 1995; Hutcheson *et al*., 2001; Pecina & Berridge, 2005; LaLumiere & Kalivas, 2008; Rogers *et al*., 2008; Bossert *et al*., 2012). GABAergic medium spiny neurons (MSNs) are the principle output neurons in the NAc and are canonically divided into subpopulations based on the expression of the dopamine receptor type 1 (D1 MSN) or D2 MSNs (Kalivas, 2009; Gerfen & Surmeier, 2011; Scofield *et al*., 2016). These two cell types show divergent roles in reinforcement (Kravitz *et al*., 2012) and respond differently to drug exposure (Kalivas, 2009; Lobo *et al*., 2010; Gerfen & Surmeier, 2011; Calipari *et al*., 2016; Scofield *et al*., 2016; van Zessen *et al*., 2021). Past work has examined plasticity in D1- and D2-MSNs produced by opioid exposure (Glass *et al*., 2008; Graziane *et al*., 2016; Hearing *et al*., 2016; Zhu *et al*., 2016; Lefevre *et al*., 2023); however, these studies have been largely input agnostic, (except Zhu et al., 2016). Further, it remains unclear whether 1) these adaptations map onto plasticity associated with more translational models of volitional drug taking, 2) plasticity is unique within mPFC-NAc circuits, and 3) adaptations are similar in females, as studies to date have been selectively in males. To address these gaps, the present study investigated sex-specific effects of self-administering remifentanil, the μOR-specific and highly potent synthetic opioid, on enduring synaptic properties of D1 and D2 MSNs in the NAc core and shell that receive afferents from the PL and IL, respectively.

## 2 MATERIALS AND METHODS

### 2.1 Animals

Adult male (N = 100; *n* = 51 – Remifentanil; *n* = 49 – Yoked Saline) and female (N = 48; *n* = 24 – Remifentanil; *n* = 24 – Yoked Saline) mice (PN45-60) were bred and raised in the animal facility at Marquette University. Animals were housed in a temperature- and humidity-controlled room maintained on a 12-h reverse light/dark cycle. BAC transgenic mice double homozygous for *Drd*1*a*−*tdTomato* and *Drd2*−*eGFP* mice (graciously provided by Dr. Rob Malenka, Stanford University) (Pascoli *et al*., 2011; Nelson *et al*., 2012) and subsequently bred with C57BL/J6 mice (Jackson Laboratories) to generate mice heterozygous for both transgenes. In some instances, mice only expressing *tdTomato* in D1 MSNs (Jackson Laboratories) were used as tdTomato signaling. For these mice, D1 MSNs were identified as tdTomato(+) and D2 MSNs were identified as tdTomato(-), an approach previously used (Hearing *et al*., 2016; Anderson *et al*., 2019). All animals were group-housed and provided with food and water *ad libitum* until the start of surgical procedures. All procedures were approved by Marquette University’s IACUC and conform to the guidelines on humane animal care.

### 2.2 Intravenous catheter surgery

Intravenous (i.v.) catheter surgeries were performed under general isoflurane (1-3%; Covetrus) anesthesia as previously described (Anderson et al., 2021). Briefly, mice were implanted with a standard mouse catheter in the right jugular vein (Access Technologies; 2/3Fr. x 6cm silicone, Cat No. AT-MJVC-2914A) and connected to a back mount (PlasticsOne; 22GA, Part No. 8I31300BM01). Following catheter surgery, mice were single-housed and allowed to recover for at least 5 days prior to beginning self-administration. Before and after self-administration sessions, catheters were immediately flushed with 0.05 mL (i.v.) of heparinized (20 IU/mL; Hospira, Inc.) bacteriostatic 0.9% saline (Hospira, Inc.) containing gentamicin sulphate (Sparhawk Laboratories, Inc.; 0.25 mg/mL) and containing enrofloxacin (Norbrook Laboratories; 4.994 mg/mL) respectively. Intravenous application of 0.05 mL of Brevital Sodium (5mg/mL) was periodically used to assess catheter patency.

### 2.3 Intracranial virus injection of ChR2-AAV

Intracranial viral surgeries were either performed the week prior to IV catheter surgeries or within 48 hours after the final day of remifentanil self-administration. pAAV9-CaMKIIa-hChR2(H134R)-EYFP was a gift from Karl Deisseroth (Addgene plasmid #26969; http://n2t.net/addgene:26969; RRID:Addgene_26969) (Lee et al., 2010). Virus was bilaterally injected (0.5 µl; 0.1 µl /min) into the PL (Males: AP: +1.8, ML: ± 0.4, DV: -2.3 mm; Females: AP: +1.75, ML: ± 0.4, DV: -2.3 mm) or IL (Males: AP: +1.8, ML: +/- 0.40, DV: -3.2mm; Females: AP: +1.75, ML: +/- 0.40, DV: -3.2mm) using standard stereotaxic apparatus attached with a 10 µL Hamilton syringe (Hamilton). Post injections, the syringe was left in place for 5 min to allow for diffusion of the virus into the tissue.

### 2.4 Self-administration

We used a ‘paired’ remifentanil self-administration paradigm that has been previously published (Anderson *et al*., 2021). Briefly, all animals were initially food restricted to 85-90% of their original weight and habituated overnight in their home cage to 25% liquid Ensure® diluted with water. On the following day, mice began training in standard operant boxes installed with a liquid dipper system (Med Associates, Inc.). Mice were trained to press the active lever which resulted in a 20 s presentation of Ensure® dipper and a simultaneous infusion of either saline (0.025 mL) or remifentanil HCl (0.025mL of 5µg/kg/infusion; Ultiva®; Mylan Institutional, LLC; purchased from Froedtert Hospital Pharmacy, Milwaukee, WI), as previously published (Anderson *et al*., 2021). Training was conducted using an increasing fixed-ratio (FR) schedule. On the first day of training, each active lever press resulted in a 20s presentation of the dipper containing Ensure® and saline/remifentanil infusion (FR1) for a maximum of 25 dipper/infusions. The following day, mice that obtained the maximum number of reinforcers underwent FR2 with a total of 50 dipper/infusion pairings. On the final day of training, 3 active lever presses were required (FR3) for each dipper/infusion presentation with a maximum of 100 reward presentations. Animals were required to reach the maximum number of reinforcers for each day (FR1 = 25, FR2 = 50, FR3 = 100) before proceeding with the next FR training. Following completion of training, mice underwent 1-3 days of forced abstinence followed by one self-administration session of the remifentanil or saline alone before food was returned *ad libitum*.

Male and female mice that successfully acquired training progressed to remifentanil/saline self-administration. Self-administration was conducted for an average of 5 days/week with each session lasting for 2-3 hours per day. During self-administration, each active lever press resulted in an i.v. infusion of remifentanil or saline (FR1) and was paired with the presentation of visual cue (light). Between infusions, a 20-second time out period was present during which levers were extended but not reinforced. A minimum of 10 remifentanil infusions and discrimination between the active and inactive lever was used as self-administration criterion. Following self-administration, mice underwent 2-3 weeks of forced abstinence in their home cage prior to electrophysiology recordings.

### 2.5 Slice electrophysiology

Two to three weeks after the last self-administration session, mice were mildly anesthetized with isoflurane (Covetrus), decapitated, the brain removed and put in oxygenated (95% O2/5% CO2) ice-cold sucrose artificial cerebrospinal fluid (sucrose ACSF; containing in mM: Sucrose 228, KCl 2.5, NaH_2_PO_4_ 1.0, MgSO_4_ 7.0, CaCl_2_ 0.5, NaHCO_3_ 26, and glucose 11) solution. Acute coronal slices (300 µm) containing the NAc core and shell were prepared with a vibratome (Leica VT1000S) and transferred to an incubation chamber filled with oxygenated ACSF (containing in mM: NaCl 119, KCl 2.5, NaH_2_PO_4_ 1.0, MgCl_2_ 1.3, CaCl_2_ 2.5, NaHCO_3_ 26.2 and glucose 11).

Slices were incubated at room temperature for 45 mins before being placed in the recording chamber. Slices were continuously superfused with oxygenated ACSF maintained at 29-32°C using a single inline solution heater (Warner Instruments) throughout recordings. MSNs were identified by their morphology and capacitance (>50 pF). Anatomical organization was used to distinguish between MSNs in NA core vs NA shell. Fluorescent tags were used to identify MSNs receiving direct input from PL or IL. When using *Drd1*a*-tdTomato* mice, D1 MSNs were identified as tdTomato(+) and putative D2 MSNs as tdTomato(-). When using *Drd1*a-*tdTomato/Drd2-eGFP* double transgenics, D1 MSNs were identified as tdTomato(+) and D2 MSNs identified as eGFP(+). Cells were checked for overlap in fluorescence in these mice to ensure recordings were from cells selectively expressing one fluorescent tag. Additionally, electrophysiological recordings were selectively conducted in MSNs located in dorsomedial NAc core and dorsal aspect of rostral NAc shell.

For miniature excitatory postsynaptic currents (mEPSCs), 0.7 mM lidocaine was added to the recording solution. mEPSCs were recorded at a holding potential (V_h_) of -72 mV with borosilicate glass pipettes (2.5-4.5 MΩ; Sutter Instruments) filled with Cs-based internal solution (containing in mM: CsMeSO_4_ 120, CsCl 15, TEA-Cl 10, NaCl 8, HEPES 10, EGTA 5, spermine 0.1, QX-314 5, ATP-Mg 4, and GTP-Na 0.3). Measurement, data collection, and analysis were performed as described (M. C. Hearing et al., 2016; Kourrich et al., 2007; Madayag et al., 2019). Series resistance was monitored continuously during all recordings, and a change beyond 20% resulted in exclusion of the cell from data analyses. Data were filtered at 2 kHz and digitized at 5 kHz via a Sutter Integrated Patch Amplifier and Igor Pro software (Wavemetrics). At the beginning of each sweep, a depolarizing step (4 mV, 100 ms) was generated to monitor series (10–40 MΩ) and input resistance (>400 MΩ). Holding potentials were corrected for liquid junction potential (∼8 mV).

AMPAR/NMDAR (A/N) ratios were computed from optically-evoked EPSCs (oEPSCs) at +40 mV with and without 50 µM D-aminophosphoonovaleric acid (D-APV; selective NMDAR antagonist). EPSCs were obtained at 0.1 Hz, with pulses of 473-nm wavelength full-field light (submerged 40x objective, Olympus) using a SOLA SE II354 Light Engine (Lumencor). Pulse duration (0.3-2-ms) and light intensity were adjusted to obtain EPSC amplitudes of ∼150-500 pA across cells. For assessment of AMPAR subunit composition, a current-voltage (I-V) curve was plotted, with AMPAR-mediated EPSCs measured at -72, +8, and +48 mV following correction of holding potential for liquid junction potential. AMPAR rectification indices were calculated by dividing the amplitude of AMPAR current at +48, +8 and -72 mV by the amplitude of AMPAR current at -72 mV. For paired-pulse ratio experiments, EPSCs were measured at 50, 100, and 200 ms interstimulus intervals (ISI). The paired-pulse ratio at individual ISI was calculated by dividing the amplitude of EPSC2:EPSC1.

### 2.6 Statistical Analysis

Statistical analyses employed mixed-, multivariate- or univariate-ANOVAs and were conducted using IBM SPSS (v30, New York NY, USA). Two separate mixed-ANOVAs were used to assess sex-specific differences in lever pressing behavior of mice that received saline or remifentanil. We also used unpaired t-test employing Welch’s correction to assess sex differences in cumulative infusions. Electrophysiology data from yoked-saline control groups was compared to assess baseline cell type- and sex-specific differences in synaptic properties of D1 vs D2 MSNs. Separate ANOVAs were conducted for analyzing baseline differences in synaptic properties of PL-Core and IL-Shell MSNs. To evaluate the influence of remifentanil treatment and sex on synaptic properties, we conducted separate ANOVAs on D1 vs D2 PL-Core MSNs. Similarly, we conducted separate ANOVAs on D1 vs D2 IL-Shell. Multivariate-ANOVA was used to assess group differences in mEPSC amplitude and frequency. Group differences in A/N ratios and AMPA rectification index were assessed using separate univariate-ANOVAs. A mixed-ANOVA was used to assess group differences in paired-pulse ratio at different ISIs. We also analyzed group differences in paired-pulse ratio at each ISI using separate univariate-ANOVAs. All ANOVAs employed Bonferroni correction for pairwise comparisons. Whenever tests of sphericity were violated, Greenhouse-Geisser corrected degrees of freedom are reported. GraphPad Prism 10.4.2 was used to graph the data.

## 3 RESULTS

To determine how contingent administration of a clinically relevant opioid, remifentanil, impacts mPFC-NAc circuit function, male and female mice underwent 10–14 days of saline or remifentanil self-administration followed by 14–22 days of forced abstinence (**Fig. 1A**). During maintenance of remifentanil self-administration (days 5-14), both males and females showed significantly greater pressing of the active lever compared to the inactive lever across all self-administration days (*Lever x Day: F*_(4.14, 302.32)_ = 5.044, *p* < 0.001), with no differences in active or inactive presses across sex (*Sex x Lever x Day interaction: F*_(4.14, 302.32)_ = 1.244, *p* = 0.292) (**Fig. 1B**). We also compared the number of remifentanil infusions between males and females during the maintenance phase of remifentanil self-administration. Number of infusions changed significantly across maintenance of remifentanil self-administration but did not differ between males and females (*Infusions*: *F*_(4.71, 343.67)_ = 4.745, *p* < 0.001; *Infusions x Sex: F*_(4.71, 343.67)_ = 1.626, *p* = 0.157) (**Fig. 1C**). Examination of cumulative infusions showed a marginally greater number of infusions in females compared to males (*t*_(45.89)_ = 2.010, *p* = 0.050) (**Fig. 1D**).

**Figure 1.**
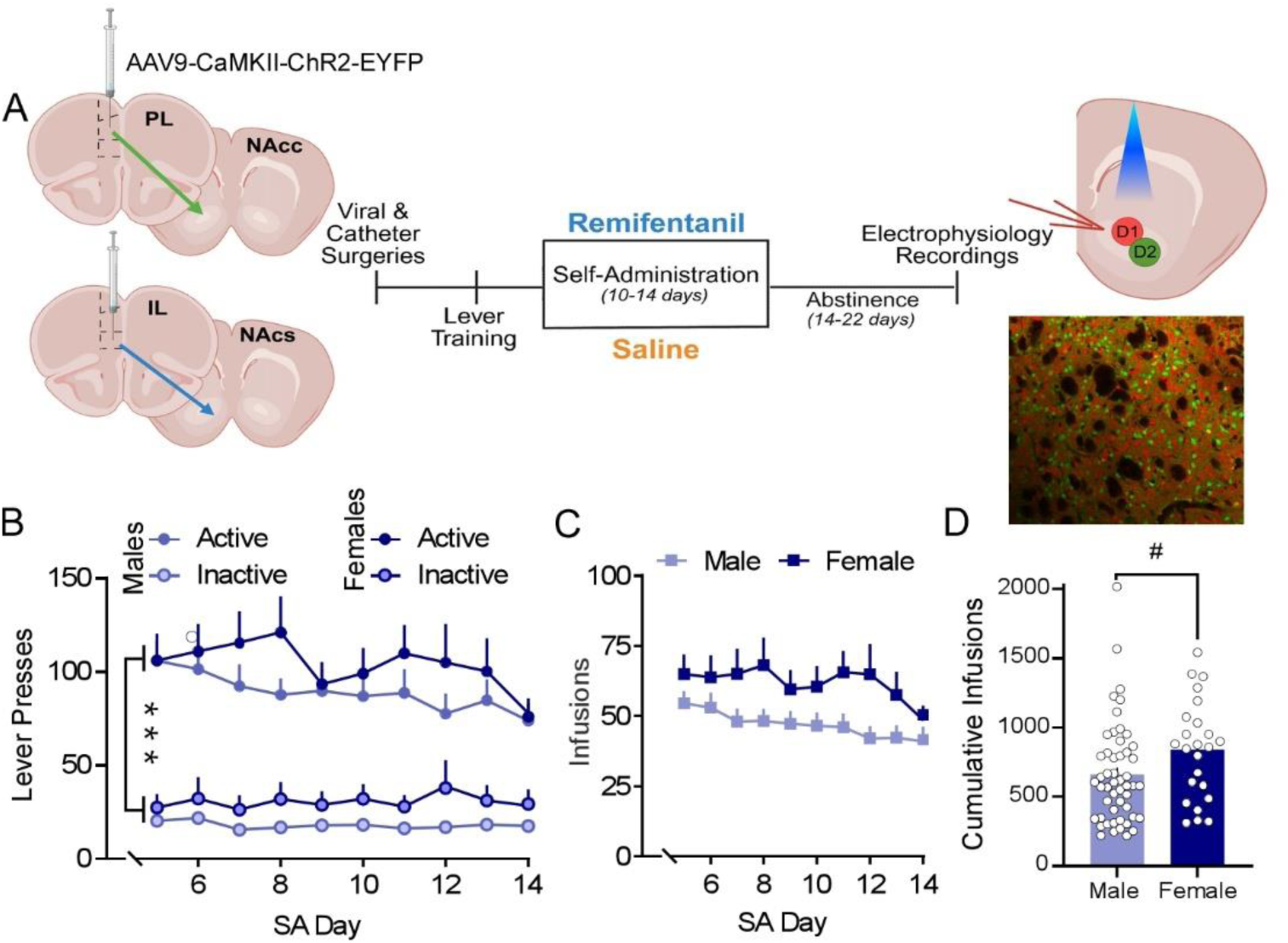
Experimental design and remifentanil self-administration in males and females. **A.** From left to right: Location of intracranial channelrhodopsin (ChR) virus (AAV9-CaMKIIa-hChR2(H134R)-eYFP) injections; self-administration, and abstinence timeline; and location of whole cell patch electrophysiology recordings. *Insert:* Representative confocal image of the NAc of *Drd*1*a*−*tdTomato/Drd*2−*eGFP* heterozygous mice expressing TdTomato (red) in D1^+^ MSNs and eGFP (green) in D2^+^ MSNs. **B**. Summary graph of lever pressing behavior during the maintenance phase (day 5-14) of remifentanil self-administration. Males (light blue circles) and females (dark blue circles) had significantly greater number of active lever presses (solid circles) vs inactive lever presses (bordered circles). **C**. Summary graph of average remifentanil infusions. Males (light blue squares) and females (dark blue squares) did not differ in daily number of remifentanil infusions. **D**. Cumulative infusions of remifentanil in females (dark blue) were marginally greater than males (light blue). Summary data are presented as Mean ± SEM. \*\*\**p <* .001 (Active vs Inactive); ^#^*p <* .05 for sex effects.

### 3.1 Sex- and cell-type specific baseline properties of NAc core and shell MSN synaptic transmission

We assessed sex- and cell-type specific baseline differences in the synaptic transmission of D1 and D2 NAc core and shell MSNs, agonistic of input, via amplitude and frequency of mEPSCs. In the NAc core, a significant sex and cell-type interaction was observed for mEPSC amplitude (*Cell-type x Sex*: *F*_(1,39)_ = 4.785, *p* = 0.035; **Fig. 2A, B**). In males, D2 MSNs exhibited greater mEPSC amplitude compared to D1 MSNs (*p* = 0.003), while the amplitude of mEPSCs in D1 MSNs were greater in females compared to males (*p* = 0.010). Examination of mEPSC frequency showed a significant effect of cell-type (*F*_(1, 39)_ = 9.964; *p* = 0.003) but not sex (*F*_(1, 39)_ = 4.006; *p* = 0.052) nor an interaction between cell-type and sex (*F*_(1,39)_ = 1.546; *p* = 0.221). Specifically, the mEPSC frequency of D2 MSNs was significantly greater than D1 MSNs (*p* = 0.003) in the NAc core (**Fig. 2C**). In the NAc shell, no significant differences in mEPSC amplitude were observed across cell-type or sex (*Cell-type*: *F*_(1,37)_ = 0.113; *p* = 0.738; *Sex*: *F*_(1,37)_ = 1.437; *p* = 0.238; *Cell-type x Sex*: *F*_(1,37)_ = 0.043; *p* = 0.837; **Fig. 2D, E**). Conversely, mEPSC frequency of NAc shell MSNs showed a significant effect of cell-type (*F*_(1,37)_ = 5.673; *p* = 0.022) with no differences observed across sex (*F*_(1,37)_ = 0.608; *p* = 0.441) or an interaction (*F*_(1,37)_ = 1.469; *p* = 0.233). Specifically, the mEPSC frequency of D2 MSNs was significantly greater than D1 MSNs (*p* = 0.022) in the NAc shell (**Fig. 2F**). Taken together, these data indicate that in males, excitatory transmission in the NAc core is greater at D2 versus D1 MSNs and that compared to males, females exhibit higher transmission at D1 MSNs.

**Figure 2.**
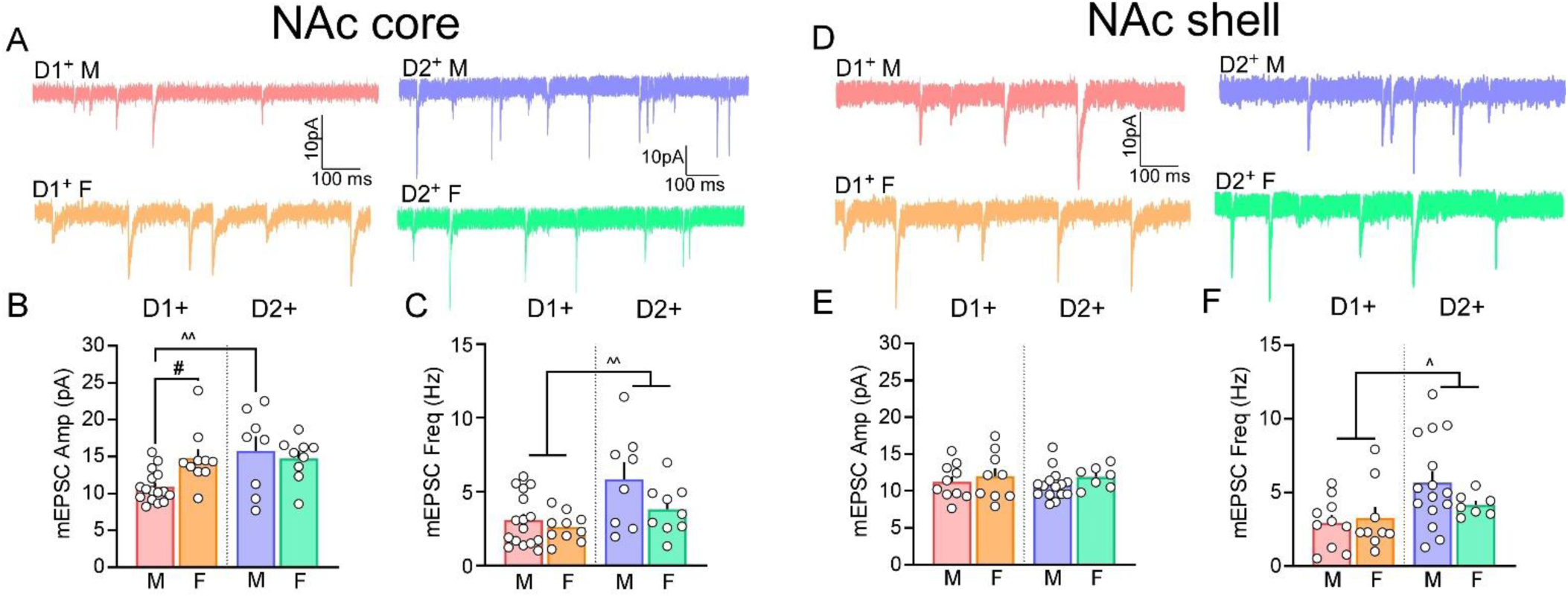
D1 and D2 MSNs of NAc core and shell show sex-specific differences in their baseline synaptic transmission. **A.** Representative traces of mEPSCs of D1 (light red) and D2 (light blue) NAc core MSNs of saline treated males (top), and D1 (light orange) and D2 (light green) NAc core MSNs of saline treated females (bottom). **B**. At baseline, mEPSC amplitude of D2 NAc core MSNs (light blue bar) is higher than D1 NAc core MSNs (light red) in males. The amplitude of mEPSCs does not differ between D1 (light orange) vs D2 (light green) NAc core MSNs of females. The amplitude of mEPSCs of females is greater than males in D1 but not D2 NAc core MSNs. **C**. The frequency of mEPSCs of D1 NAc core MSNs of males (light red) and females (light blue) is lower than mEPSC frequency of D2 NAc core MSNs of males (light blue) and females (light green). **D**. Representative traces of mEPSCs of D1 (light red) and D2 (light blue) NAc shell MSNs of saline treated males (top), and D1 (light orange) and D2 (light green) NAc shell MSNs of saline treated females (bottom). **E**. There were no differences in the mEPSC amplitude of D1 (light red) or D2 (light blue) NAc shell MSNs of males, and D1 (light orange) or D2 (light green) NAc shell MSNs of females. **F**. The frequency of mEPSCs of D1 NAc core MSNs of males (light red) and females (light blue) is lower than mEPSC frequency of D2 NAc core MSNs of males (light blue) and females (light green). D1^+^/D2^+^ - D1/D2-expressing MSNs; M – males, F – females; Summary data are presented as Mean ± SEM. ^, ^^*p <* 0.05, 0.01 for cell-type differences; ^#^*p* < .05 for sex effects.

### 3.2 Sex- and cell-type specific baseline properties of PL-Core and IL-Shell MSN synaptic transmission

We next examined whether cell-type or sex-specific differences in baseline pre- and postsynaptic signaling were present at isolated PL-Core and IL-Shell synapses using oEPSCs. Examination of A/N ratio in PL-Core circuits showed a significant effect of cell-type (*F*_(1, 26)_ = 31.431, *p* < 0.001) but not sex (*F*_(1, 26)_ = 1.890, *p* = 0.181), or an interaction between cell-type and sex (*F*_(1, 26)_ = 0.097, *p* = 0.758). Specifically, A/N ratios were greater at D1 compared to D2 PL-Core synapses (*p* < 0.001; **Fig. 3A, B**). To test whether these differences reflected distinctions in AMPAR subunit composition, we evaluated current-voltage (I-V) relationships of oEPSCs. A significant effect of cell-type (*F*_(1, 26)_ = 10.284, *p* = 0.004) but no effect of sex (*F*_(1, 26)_ = 1.809, *p* = 0.190) or interaction between cell-type and sex (*F*_(1, 26)_ = 2.050, *p* = 0.164) was observed, with AMPAR rectification indices lower in D2 versus D1 PL-Core synapses (*p* = 0.004; **Fig. 3C-E**). These differences in the AMPAR rectification indices indicate that lower A/N ratios observed at D2 PL-Core synapses likely reflect an increased presence of calcium-permeable AMPARs (CP-AMPARs) at baseline. Examination of presynaptic release probability using paired-pulse ratios showed that paired-pulse ratios significantly changed at increasing ISIs (*F*_(1.25, 42.36)_ = 24.755, *p* < 0.001). But there was no significant effect of cell-type (*F*_(1.25, 42.36)_ = 1.539, *p* = 0.222), sex (*F*_(1.25, 42.36)_ = 0.497, *p* = 0.525) or an interaction between cell-type and sex (*F*_(1.25, 42.36)_ = 0.318, *p* = 0.625). However, because shorter ISI durations are more often shaped by residual presynaptic Ca2+ while longer ISIs may reflect recover kinetics at presynaptic sites (Dittman *et al*., 2000), we conducted subsequent analyses in paired-pulse ratio at 50, 100 and 200 ms ISI separately. At 50 ms ISI, we found a significant effect of cell-type (*F*_(1, 34)_ = 11.297, *p* = 0.002) and sex (*F*_(1, 34)_ = 6.980, *p* = 0.012) but did not see a significant interaction between cell- type and sex (*F*_(1, 34)_ = 0.381, *p* = 0.541). At 100 ms ISI, we found a significant effect of cell-type (*F*_(1, 34)_ = 4.224, *p* = 0.048) and sex (*F*_(1, 34)_ = 6.892, *p* = 0.013) but did not see a significant interaction between cell-type and sex (*F*_(1, 34)_ = 1.501, *p* = 0.229). At 200 ms ISI, we found a significant effect of sex (*F*_(1, 34)_ = 8.164, *p* = 0.007), but did not see an effect of cell-type (*F*_(1, 34)_ = 2.711, *p* = 0.109) nor a significant interaction between cell-type and sex (*F*_(1, 34)_ = 0.893, *p* = 0.351; **Fig. 3F, G**). Taken together, these results indicate that the decrease in presynaptic release probability at D1 vs D2 PL-Core synapses is likely driven by males while not differing in females.

**Figure 3.**
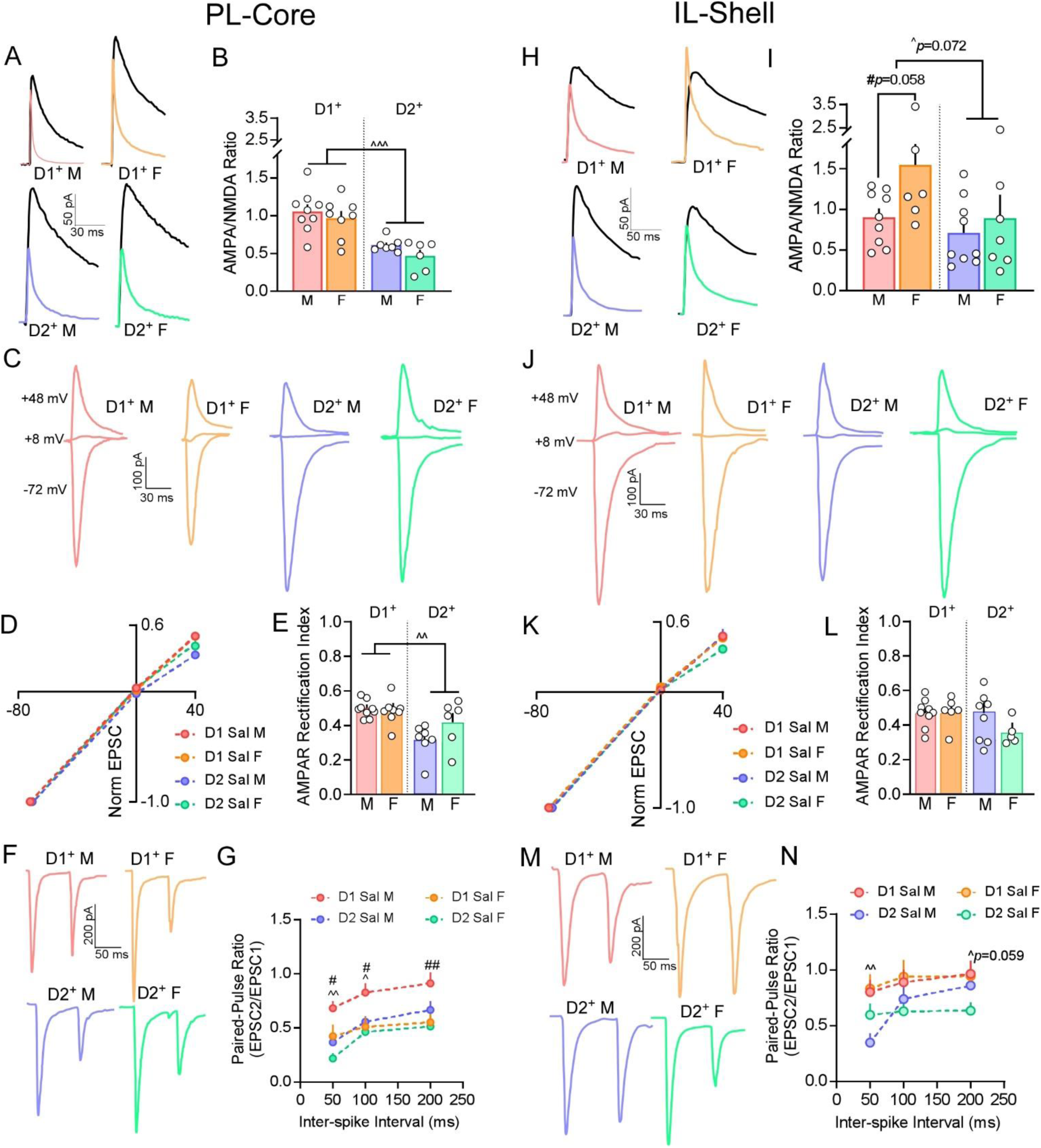
Sex- and cell-type specific differences in baseline pre- and postsynaptic signaling at PL-Core and IL-Shell synapses. **A.** Representative baseline traces of AMPAR and NMDAR mediated currents at PL-Core synapses. NMDAR current (black) was measured at +40 mV and AMPAR current (colored) was measured at +40 mV in the presence of D-APV. **B**. Summary graphs of AMPA/NMDA (A/N) ratios showing that at baseline, D1 PL-Core synapses in both males (light red) and females (light orange) exhibited significantly higher A/N ratios compared to D2 PL-Core synapses of males (light blue) and females (light green), with no effect of sex. **C**. Representative baseline oEPSC traces at -72, +8, and +48 mV at D1 and D2 PL-Core synapses in males and females. **D**. Baseline I-V relationships representing AMPAR rectification curve at D1 and D2 PL-Core synapses. **E**. Summary graph of AMPAR rectification index at +48 mV. At baseline, AMPAR rectification was significantly lower at D2 PL-Core synapses of males (light blue) and females (light green) vs. D1 PL-Core synapses of males (light red) and females (light orange). **F**. Representative traces of paired oEPSCs at 50 ms ISI at baseline in D1 and D2 PL-Core synapses. **G**. Summary graph of baseline paired-pulse ratios at 50, 100 and 200 ms ISI. At 50 and 100 ms ISI, paired-pulse ratio at D1 PL-Core synapses were higher than that at D2 PL-Core synapses. D1 PL-Core synapses of males also had higher PPRs compared to females at all ISIs. **H.** Representative traces of A/N ratios at IL-Shell synapses. NMDA current (black) was measured at +40 mV and AMPA current (colored) was measured at +40 mV in the presence of D-APV. **I**. Summary graphs of A/N ratios showing a trend toward increased A/N ratios was observed at D1 vs. D2 IL-Shell synapses, potentially driven by higher ratios in females. **J**. Representative oEPSC traces at -72, +8, and +48 mV at D1 and D2 IL-Shell synapses. **K**. I-V relationships representing AMPAR rectification curve at D1 and D2 IL-Shell synapses. **L**. Summary graph of AMPAR rectification index at +48 mV at D1 vs D2 IL-Shell synapses showing no differences in rectification. **F**. Representative traces of paired pulses evoked with 50 ms ISI at D1 and D2 IL-Shell synapses. **G**. Summary graph of paired-pulse ratios at 50, 100 and 200 ms ISI. At 50 ms ISI, paired-pulse ratio at D1 IL-Shell synapses were higher than that at D2 IL-Shell synapses while at 200 ms ISI, they were marginally higher. D1^+^/D2^+^ - D1/D2-expressing MSNs, M – males, F – females; Summary data are presented as Mean ± SEM. ^, ^^, ^^^*p <* 0.05, 0.01, 0.001 for cell-type differences; ^#^, ^##^*p* < 0.05, 0.01 for sex effects.

Examination of IL-Shell inputs showed distinctly different patterns of baseline signaling. While not significant, a cell-type (*F*_(1, 27)_ = 3.507, *p* = 0.072) and sex-specific (*F*_(1, 27)_ = 3.346, *p* = 0.078) trend towards increased A/N ratio was observed at D1 vs D2 IL-Shell synapses (*p* = 0.072) likely driven by increased A/N ratio at D1 IL-Shell synapses of females vs males (*p* = 0.058; **Fig. 3H, I**). AMPAR rectification indices at D1 and D2 IL-Shell synapses did not show significant differences across cell-type (*F*_(1, 25)_ = 1.007, *p* = 0.325), sex (*F*_(1, 25)_ = 1.737, *p* = 0.199), nor was there an interaction between cell-type and sex (*F*_(1, 25)_ = 1.007, *p* = 0.325; **Fig. 3J-L**). In contrast, assessment of release probability via paired-pulse ratios showed a significant interaction of cell-type and sex across all ISIs (*F*_(2, 66)_ = 7.861, *p* < 0.001). *Post hoc* pairwise comparisons revealed that within sex, paired-pulse ratios at male D1 IL-Shell synapses were higher at the 50 ms ISI compared to D2 IL-Shell synapses (*p* = 0.006), whereas ratios at female D1 IL-Shell synapses were marginally higher at the 200 ms ISI compared D2 IL-Shell synapses (*p* = 0.059; **Fig. 3M, N**).

### 3.3 Cell-type specific effects of remifentanil on D1 and D2 MSN signaling in NAc core and shell

Previous work has shown that non-contingent opioid exposure promotes cell-type and region-specific adaptations in the NAc (Hearing *et al*., 2016; Madayag *et al*., 2019); however, it remains unclear whether similar forms of pathway agnostic plasticity are observed after volitional opioid taking and whether this plasticity is isolated to select pathways within mPFC-NAc circuits.

Initial examination of agnostic synaptic transmission (assessed by measuring mEPSCs) showed a significant interaction of treatment (saline, remifentanil) and sex (male, female) in the amplitude of mEPSCs of D1 NAc core MSNs (*F*_(1, 56)_ = 5.485, *p* = 0.023). Within sex, remifentanil males (*p* < 0.001) but not females (*p* = 0.573) showed significantly greater mEPSC amplitudes compared to saline counterparts. Additionally, within treatment groups, mean amplitude of mEPSCs of D1 NAc core MSNs was significantly greater in saline females compared to saline males (*p* = 0.003), however no difference was observed in remifentanil treated mice (*p* = 0.914; **Fig. 4A, B**). Assessment of mEPSC frequency in D1 NAc core MSNs showed a significant effect of treatment (*F*_(1, 56)_ = 5.780, *p* = 0.020) in that remifentanil increased mEPSC frequency. However, no significant effect of sex (*F*_(1, 56)_ = 1.604, *p* = 0.211) or a significant interaction between treatment and sex (*F*_(1, 56)_ = 0.143, *p* = 0.706; **Fig. 4C**) was observed. In D2 NAc core MSNs, a main effect of treatment was observed for both mEPSC amplitude (*F*_(1, 32)_ = 6.392, *p* = 0.017) and frequency (*F*_(1, 32)_ = 16.886; *p* < 0.001), with remifentanil reducing both, compared to saline controls. No significant effect of sex (*amplitude: F*_(1, 32)_ = 0.095, *p* = 0.760); *frequency:* (*F*_(1, 32)_ = 2.57, *p* = 0.119) or a significant interaction between treatment and sex (*amplitude: F*_(1, 32)_ = 1.231; *p* = 0.276; *frequency*: *F*_(1, 32)_ = 2.742, *p* = 0.108) was observed (**Fig. 4D-F)**. Taken together, these data indicate that remifentanil upregulates excitatory signaling at D1 NAc core MSNs primarily in males, whereas there is a general reduction at D2 NAc core MSNs in both sexes.

**Figure 4.**
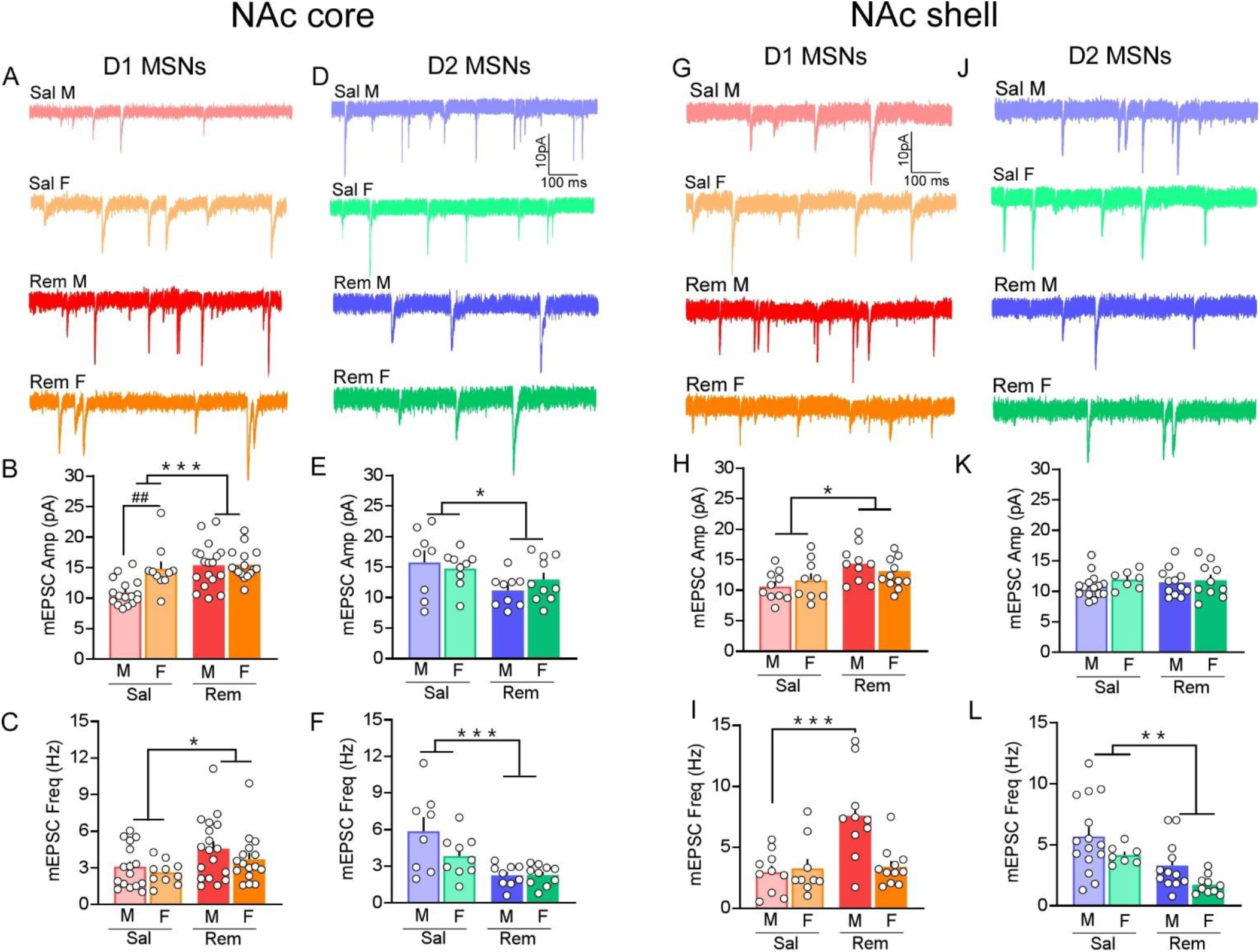
Divergent sex- and cell-type specific changes in synaptic transmission of NAc core and shell MSNs are induced after remifentanil treatment. **A.** Representative mEPSC traces of D1 NAc core MSNs. **B**. Summary graph of mEPSC amplitude in D1 NAc core MSNs demonstrating that remifentanil treatment selectively increased mEPSC amplitude in males (red bars) and females (orange bars), while saline-treated females (light orange bars) exhibited higher baseline amplitudes compared to saline males (light red bars). **C**. Summary graph of mEPSC frequency in D1 NAc core MSNs showing that the frequency of mEPSCs was increased after remifentanil treatment across both sexes. **D**. Representative mEPSC traces of D2 NAc core MSNs. **E, F**. In D2 NAc core MSNs remifentanil treatment selectively decreased amplitude (**E**) and frequency (**F**) of mEPSCs in males (blue bars) and females (green bars) compared to saline treated males (light blue) and females (light green). **G**. Representative mEPSC traces of D1 NAc shell MSNs. **H**. Summary graph of mEPSC amplitude in D1 NAc shell MSNs demonstrating that remifentanil treatment selectively increased mEPSC amplitude in males and females. **I**. Summary graph of mEPSC frequency in D1 NAc shell MSNs showing that the frequency of mEPSCs was increased after remifentanil treatment in males but not females. **J**. Representative mEPSC traces of D2 NAc shell MSNs. **K**. Summary graph in D2 NAc shell MSNs showing that remifentanil treatment did not affect the amplitude of mEPSCs in both males and females. **L**. Summary graph in D2 NAc shell MSNs demonstrating that remifentanil treatment decreased mEPSC frequency in males (blue bars) and females (green bars) compared to saline treated males (light blue) and females (light green). Sal – Saline, Rem – Remifentanil, M – males, F – females; Summary data are presented as Mean ± SEM. *, **, \*\*\**p* < 0.05, 0.01, 0.001 for treatment effects; ^##^*p* < 0.01 for sex effects.

To assess pathway agnostic plasticity in D1 and D2 NAc shell MSNs, we measured remifentanil-induced alterations in the amplitude and frequency of mEPSCs. Remifentanil significantly increased the amplitude of mEPSCs of D1 NAc shell MSNs (*F*_(1, 36)_ = 4.368, *p* = 0.044). However, there was no significant effect of sex (*F*_(1, 36)_ = 0.305, *p* = 0.584) or a significant interaction between treatment and sex (*F*_(1, 36)_ = 2.005, *p* = 0.165; **Fig4. G, H**). The frequency of mEPSCs of D1 NAc shell MSNs showed a significant interaction between treatment and sex (*F*_(1, 36)_ = 8.598, *p* = 0.006). Specifically, mEPSC frequency increased significantly in D1 NAc shell MSNs of males but not females (*p* < 0.001) after remifentanil treatment vs saline (**Fig. 4I**). In D2 NAc shell MSNs, there was neither a significant effect of remifentanil treatment on the amplitude of mEPSCs (*F*_(1, 40)_ = 0.141, *p* = 0.709) across sex (*F*_(1, 40)_ = 1.049, *p* = 0.312) nor was there a significant interaction between treatment and sex (*F*_(1, 40)_ = 0.271, *p* = 0.606; **Fig. 4J, K**). However, remifentanil treatment significantly decreased the frequency of D2 NAc shell MSNs (*F*_(1, 40)_ = 12.865, *p* < 0.001) in both males and females (*F*_(1, 40)_ = 5.304, *p* = 0.027) but no significant interaction between treatment and sex (*F*_(1, 40)_ = 0.003, *p* = 0.958; **Fig. 4L**) was observed.

### 3.4 Impact of remifentanil on PL-Core D1 and D2 MSN pre- and postsynaptic signaling

Past work has examined cell-type specific plasticity following non-contingent opioid exposure (Graziane *et al*., 2016; Hearing *et al*., 2016) or assessed plasticity within select circuits including inputs from the IL to D1 MSNs (Hearing *et al*., 2016) and paraventricular nucleus of the thalamus (Zhu *et al*., 2016) input to D1 and D2 MSNs. However, no studies to date have compared plasticity associated with volitional opioid taking or directly compared plasticity within sub-circuits and cell-types. Given the roles of PL and IL subregions in opioid-related behavior, we next examined whether inputs from these cortical regions to NAc MSNs differs following remifentanil and abstinence.

Examination of A/N ratios at D1 PL-Core synapses showed a significant effect of treatment, with remifentanil reducing A/N ratios in both males and females compared to their saline counterparts (*F*_(1, 32)_ = 41.864, *p* < 0.001). No significant effect of sex (*F*_(1, 32)_ = 0.623, *p* = 0.436) or interaction between sex and treatment (*F*_(1, 32)_ = 0.137, *p* = 0.714) was observed (**Fig. 5A, B**). Assessment of AMPAR rectification showed a significant reduction in indices in remifentanil male and female mice compared to saline controls (*F*_(1, 30)_ = 33.057, *p* < 0.001).

**Figure 5.**
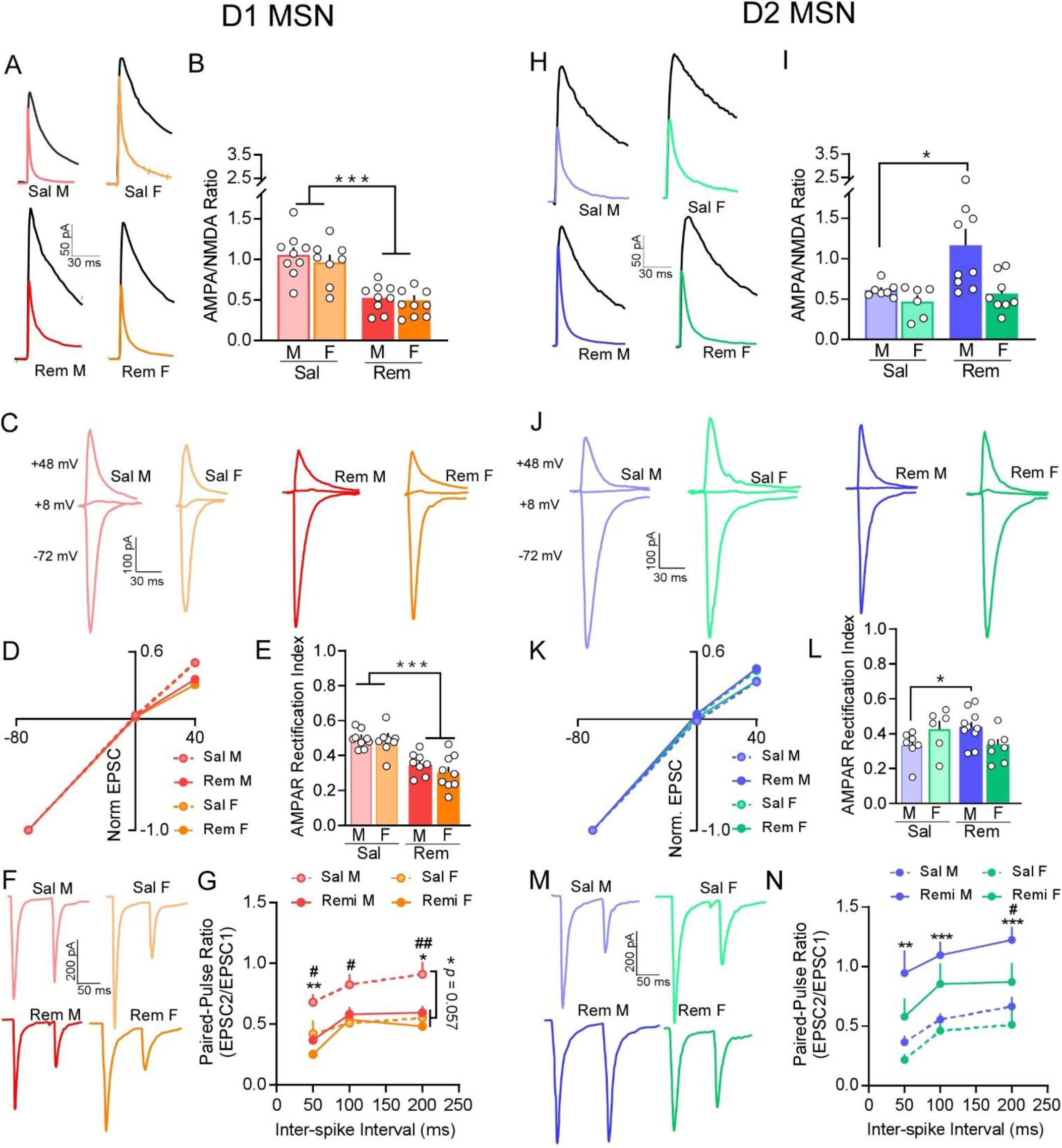
Pre- and postsynaptic signaling mechanisms after remifentanil treatment differ between D1 and D2 PL-Core synapses. **A.** Representative traces of AMPAR and NMDAR mediated currents at D1 PL-Core synapses in saline (top) vs remifentanil treated (bottom) males (left) and females (right). NMDAR current (black) was measured at +40 mV and AMPAR current (colored) was measured at +40 mV in the presence of D-APV. **B**. Summary graph of A/N ratios shows that remifentanil significantly reduced A/N ratios at D1 PL-Core synapses in both males and females compared to saline controls. **C**. Representative oEPSC traces at -72, +8, and +48 mV at D1 PL-Core synapses in saline and remifentanil treated males and females. **D**. I-V plots derived from oEPSC traces show decreased rectification following remifentanil at D1 PL-Core synapses in both males and females. **E**. Summary graph of AMPAR rectification index at +48 mV confirmed that remifentanil significantly decreased the AMPAR rectification index at D1 PL-Core synapses across both sexes. **F**. Representative paired oEPSC traces at 50 ms ISI at D1 PL-Core synapses in saline (top) and remifentanil (bottom) treated males (left) and females (right). **G**. Summary graph of paired-pulse ratios at 50, 100 and 200 ms ISI in D1 PL-Core synapses demonstrated that remifentanil marginally decreased the paired-pulse ratio. Specifically, paired-pulse ratio was significantly decreased after remifentanil treatment at 50, and 200 ms ISI. Paired-pulse ratios were significantly decreased after remifentanil treatment in males but not females at all ISIs tested. **H.** Representative traces of AMPAR and NMDAR mediated currents at D2 PL-Core synapses in saline (top) and remifentanil treated (bottom) males (left) and females (right). NMDAR current (black) was measured at +40 mV and AMPAR current (colored) was measured at +40 mV in the presence of D-APV. **I**. Summary graph of A/N ratios shows that remifentanil significantly increased A/N ratios at D2 PL-Core synapses in males but not females compared to saline controls. **J**. Representative oEPSC traces at -72, +8, and +48 mV at D2 PL-Core synapses in saline vs remifentanil treated males and females. **K**. I-V plots derived from oEPSC traces show increased rectification following remifentanil treatment at D2 PL-Core synapses in males but not females. **L**. Summary graph of AMPAR rectification index at +48 mV confirmed that remifentanil significantly increased the AMPAR rectification index at D2 PL-Core synapses of males. **M**. Representative paired oEPSC traces at a 50 ms ISI at D2 PL-Core synapses in saline (top) vs. remifentanil (bottom) treated males (left) and females (right). **G**. Summary graph of paired-pulse ratios at 50, 100 and 200 ms ISI in D2 PL-Core synapses demonstrated that remifentanil significantly increased the paired-pulse ratios at all ISI tested. At 200 ms ISI paired-pulse ratio of males was significantly higher than females. Sal – Saline, Rem – Remifentanil, M – males, F – females; Summary data are presented as Mean ± SEM. *, **, \*\*\**p* < 0.05, 0.01, 0.001 for treatment effects; ^#^, ^##^*p* < 0.05, 0.01 for sex effects.

However, we did not observe a significant effect of sex (*F*_(1, 30)_ = 0.741, *p* = 0.396) or a significant interaction between sex and treatment (*F*_(1, 30)_ = 0.568, *p* = 0.457; **Fig. 5C-E)**. Examination of release probability showed a marginally significant impact of remifentanil treatment on paired-pulse ratio at D1 PL-Core synapses (*ISI x Treatment*: *F*_(1.31, 52.28)_ = 3.485, *p* = 0.057). However, we did not observe any significant effect of sex (*ISI x Sex*: *F*_(1.31, 52.28)_ = 0.731, *p* = 0.484) or an interaction between sex and treatment on paired-pulse ratio (*ISI x Treatment x Sex*: *F*_(1.31, 52.28)_ = 0.925, *p* = 0.366). We also assessed changes in paired-pulse ratio of D1 PL-Core synapses at 50, 100 and 200 ms ISI separately. At 50 ms ISI, we found a significant effect of remifentanil treatment (*F*_(1, 40)_ = 11.790, *p* = 0.001) on paired-pulse ratio of D1 PL-Core synapses such that remifentanil treatment decreased paired-pulse ratio vs saline treatment. We also found a significant effect of sex (*F*_(1, 40)_ = 7.073, *p* = 0.011) with paired-pulse ratio at D1 PL-Core synapses of males being greater than females. However, the interaction between treatment and sex (*F*_(1, 40)_ = 1.011, *p* = 0.321) was not significant. At 100 ms ISI, we found a significant effect of sex (*F*_(1, 40)_ = 5.858, *p* = 0.020) and marginally significant interaction between treatment and sex (*F*_(1, 40)_ = 3.346, *p* = 0.075), but no significant effect of remifentanil treatment (*F*_(1, 40)_ = 2.118, *p* = 0.153). Specifically, remifentanil treatment significantly decreased paired-pulse ratio at D1 PL-Core synapses of males vs saline (*p* = 0.024) but not in females (*p* = 0.795). Within saline treated mice, however, paired-pulse ratio at D1 PL-Core synapses of males was significantly greater than that of saline treated females (*p* = 0.006). At 200 ms ISI, we found a significant effect of treatment (*F*_(1, 40)_ = 6.531, *p* = 0.015), with the paired-pulse ratio at D1 PL-NAc synapses decreasing significantly after remifentanil treatment vs saline (*p* = 0.015). We also observed a significant effect of sex (*F*_(1, 40)_ = 9.826, *p* = 0.003) with paired-pulse ratio at D1 PL-Core synapses being greater for males vs females. However, we did not see a significant interaction between treatment and sex (*F*_(1, 40)_ = 2.694, *p* = 0.109; **Fig. 5F, G**). These data indicate that remifentanil increased postsynaptic strength at D1 PL-Core synapses across sex through upregulation of CP-AMPA receptors, whereas increases in presynaptic strength are specific to males, due in part to higher release under drug naïve conditions in females.

In contrast to D1 MSNs, a significant effect of remifentanil treatment (*F*_(1, 26)_ = 5.687, *p* = 0.025) and sex (*F*_(1, 26)_ = 7.245, *p* = 0.012) was observed with A/N ratios at D2 PL-Core synapses. While not significant, a trend towards an interaction between sex and treatment was observed (*F*_(1, 26)_ = 3.604, *p* = 0.069). Assessment of pairwise comparisons demonstrated that a significant effect of remifentanil treatment was driven by an increase in A/N ratios at D2 PL-Core synapses of remifentanil treated males compared to saline controls (*p* = 0.006; **Fig. 5H, I**). Examination of AMPAR rectification indices of D2 PL-Core synapses identified an interaction between treatment and sex (*F*_(1, 27)_ = 6.821, *p* = 0.015), with indices at D2 PL-Core synapses of males significantly greater in remifentanil treated mice vs saline controls (*p* = 0.033). No significant differences were observed at D2 PL-Core synapses between male vs females within saline (*p* = 0.089) or remifentanil groups (*p* = 0.061), nor was and difference observed across treatment in females (*p* = 0.150; **Fig. 5J-L**). Examination of release probability at D2 PL-Core synapses did not demonstrate an overall significant effect of remifentanil treatment on paired-pulse ratio across ISIs (*F*_(1.25, 41.20)_ = 0.001, *p* = 0.986) nor a significant effect of sex (*F*_(1.25, 41.20)_ = 0.621, *p* = 0.469). There was also no significant interaction between treatment and sex across ISIs (*F*_(1.25, 41.20)_ = 0.098, *p* = 0.811). However, examinations of changes in paired-pulse ratio of D2 PL-Core synapses at 50, 100 and 200 ms ISI separately demonstrated significant group differences. At 50 ms ISI, we found a significant effect of remifentanil treatment (*F*_(1, 33)_ = 11.902, *p* = 0.002) and a marginally significant effect of sex (*F*_(1, 33)_ = 3.634, *p* = 0.065) on paired-pulse ratio of D2 PL-Core MSNs. Specifically, paired-pulse ratio was significantly increased at D2 PL-Core synapses after remifentanil treatment vs saline, and it was significantly greater in males vs females (*p* = 0.065). However, the interaction between treatment and sex (*F*_(1, 33)_ = 0.559, *p* = 0.460) was not significant. At 100 ms ISI, we found a significant effect of treatment (*F*_(1, 33)_ = 17.602, *p* < 0.001) in that paired-pulse ratio was increased at D2 PL-Core synapses after remifentanil treatment vs saline. However, we did not observe a significant effect of sex (*F*_(1, 33)_ = 2.476, *p* = 0.125) or a significant interaction between treatment and sex (*F*_(1, 33)_ = 0.313, *p* = 0.579) at these synapses. At 200 ms ISI, we found a significant effect of treatment (*F*_(1, 33)_ = 15.755, *p* < 0.001) with paired-pulse ratio at D2 PL-Core synapses increasing significantly after remifentanil treatment vs saline. We also observed a significant effect of sex (*F*_(1, 33)_ = 5.059, *p* = 0.031) with paired-pulse ratio at D2 PL-Core synapses being greater for males vs females. However, we did not see a significant interaction between treatment and sex (*F*_(1, 33)_ = 0.525, *p* = 0.474; **Fig. 5M, N**). Together these data indicated that in males but not females, remifentanil alters the composition of AMPAR subunits through downregulation of CP-AMPA receptors, while presynaptic strength is increased.

### 3.5 Impact of remifentanil on IL-Shell D1 and D2 MSN pre- and postsynaptic signaling

Previous work has shown that non-contingent morphine exposure promotes pathway non-specific adaptations in NAc shell MSNs and that adaptations at D1 IL-Shell synapses play a role in opioid reward behavior (Hearing *et al*., 2016). Therefore, we next examined whether remifentanil produced overlapping or divergent forms of plasticity at IL-Shell synapses. At D1 IL-Shell synapses, mean A/N ratios were marginally increased by remifentanil treatment (*F*_(1, 31)_ = 3.925, *p* = 0.056) across sex. There was also no significant effect of sex (*F*_(1, 31)_ = 2.499, *p* = 0.124), but we found a *marginally* significant interaction between sex x treatment (*F*_(1, 31)_ = 3.966, *p* = 0.055). Assessment of pairwise comparisons demonstrated that while A/N ratio did not differ between males vs females (*p* = 0.760) after remifentanil treatment, it was decreased at D1 IL-Shell synapses of females vs saline control (*p =* 0.017; **Fig. 6A, B**). Assessment of AMPAR rectification index showed a significant treatment effect, with indices lower in remifentanil treated mice compared to saline controls (*F*_(1, 30)_ = 5.584, *p* = 0.025). However, there was no significant effect of sex (*F*_(1, 23)_ = 0.234, *p* = 0.632) or an interaction effect between sex and treatment (*F*_(1, 30)_ = 0.857, *p* = 0.362; **Fig. 6C-E**). Examination of paired-pulse ratios showed that release probability remained unchanged by remifentanil treatment across ISIs (*ISI x Treatment*: *F*_(1.54, 64.70)_ = 0.066, *p* = 0.894), with no impact of sex (*ISI x Sex*: *F*_(1.54, 64.70)_ = 0.211, *p* = 0.752) nor a significant interaction between treatment and sex (*ISI x Treatment x Sex*: *F*_(1.54, 64.70)_ = 0.395, *p* = 0.621) at D1 IL-Shell synapses. However, subsequent comparisons within 50, 100 and 200 ms ISIs separately demonstrated significant alterations. Specifically, we found a significant reduction in paired-pulse ratio at D1 IL-Shell synapses of remifentanil versus saline controls at 50 ms ISI (*F*_(1, 42)_ = 4.582, *p* = 0.038), with trends towards a reduction at 100 ms (*Treatment: F*_(1, 42)_ = 3.145, *p* = 0.083) and 200 ms (*F*_(1, 42)_ = 3.860, *p* = 0.056). However, no significant effect of sex (50 ms: *F*_(1, 42)_ = 0.665, *p* = 0.419; 100 ms: *F*_(1, 42)_ = 0.308, *p* = 0.582; 200 ms: *F*_(1, 42)_ = 0.165, *p* = 0.687) or a significant interaction between treatment and sex (50 ms: *F*_(1, 42)_ = 0.299, *p* = 0.587; 100 ms: *F*_(1, 42)_ = 0.018, *p* = 0.893; 200 ms: *F*_(1, 42)_ = 0.320, *p* = 0.575) was observed at any ISI at these synapses (**Fig. 6F, G)**.

**Figure 6.**
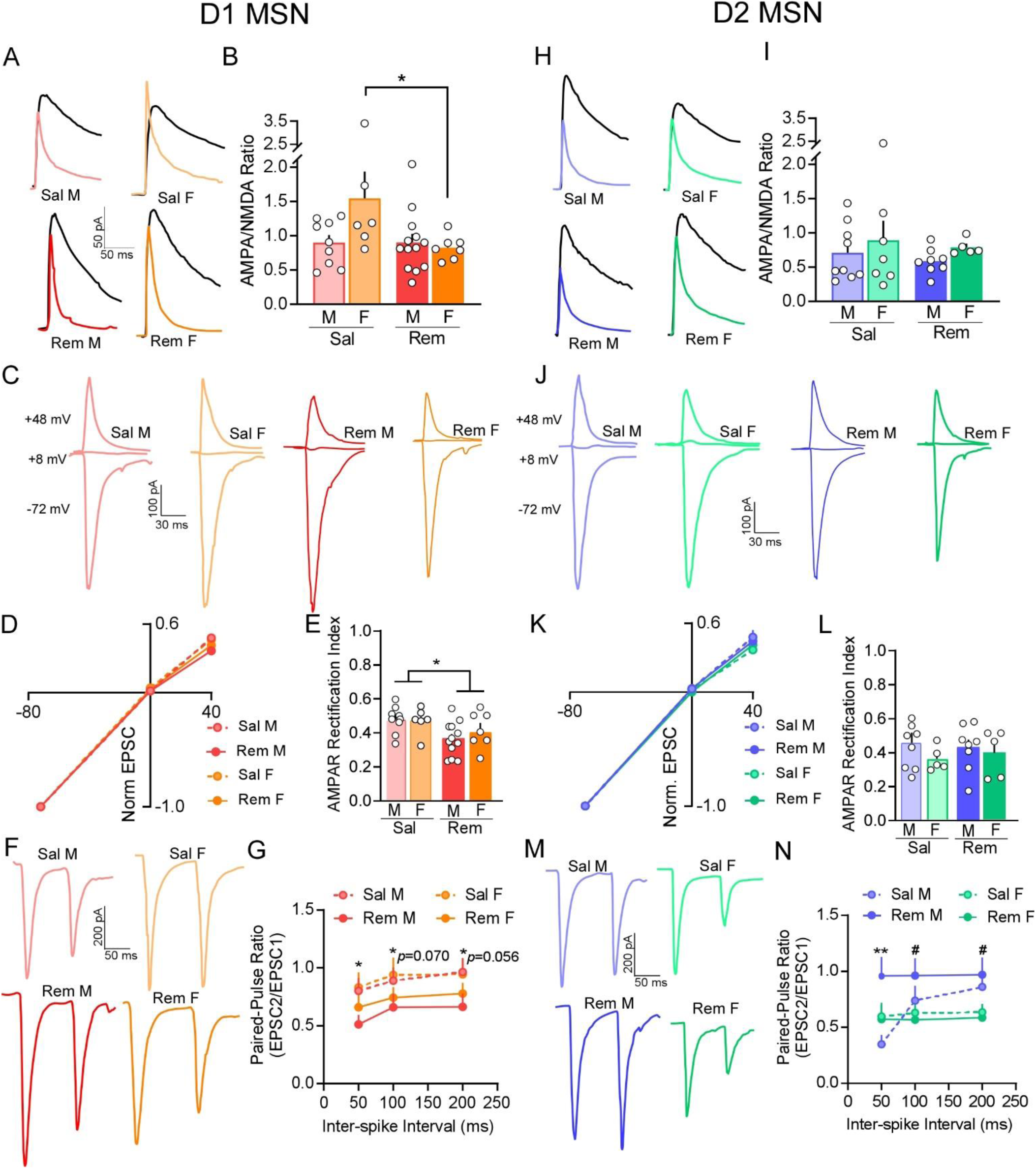
Divergent sex-specific presynaptic signaling mechanisms are induced in D1 and D2 IL-Shell synapses after remifentanil treatment. **A.** Representative traces of AMPAR and NMDAR mediated currents at D1 IL-Shell synapses in saline (top) vs remifentanil treated (bottom) males (left) and females (right). NMDAR current (black) was measured at +40 mV and AMPAR current (colored) was measured at +40 mV in the presence of D-APV. **B**. Summary graph of A/N ratios shows that remifentanil significantly reduced A/N ratios at D1 IL-Shell synapses in females but not males compared to saline controls. **C**. Representative oEPSC traces at -72, +8, and +48 mV at D1 IL-Shell synapses of saline vs remifentanil treated males and females. **D**. I-V plots derived from oEPSC traces show that remifentanil did not change AMPAR rectification at D1 IL-Shell synapses in both males and females. **E**. However, the summary graph of AMPAR rectification index at +48 mV demonstrates that remifentanil significantly decreased the AMPAR rectification index at D1 IL-Shell synapses across both sexes. **F**. Representative paired oEPSC traces at a 50 ms ISI at D1 IL-Shell synapses in saline (top) vs remifentanil (bottom) treated males (left) and females (right). **G**. Summary graph of paired-pulse ratios at 50, 100 and 200 ms ISI in D1 IL-Shell synapses demonstrated that remifentanil marginally decreased the paired-pulse ratio across sexes at all ISIs tested. **H.** Representative traces of AMPAR and NMDAR mediated currents at D2 IL-Shell synapses of saline (top) and remifentanil treated (bottom) males (left) and females (right). NMDAR current (black) was measured at +40 mV and AMPAR current (colored) was measured at +40 mV in the presence of D-APV. **I**. Summary graph of A/N ratios shows that remifentanil did not alter A/N ratios at D2 IL-Shell synapses. **J**. Representative oEPSC traces at -72, +8, and +48 mV at D2 IL-Shell synapses of saline and remifentanil treated males and females. **K**. I-V plots derived from oEPSC traces show no differences in rectification following remifentanil treatment at D2 IL-Shell synapses. **L**. Summary graph of AMPAR rectification index at +48 mV confirmed that remifentanil did not alter AMPAR rectification index at D2 IL-Shell synapses. **M**. Representative paired oEPSC traces at a 50 ms ISI at D2 IL-Shell synapses in saline (top) and remifentanil (bottom) treated males (left) and females (right). **G**. Summary graph of paired-pulse ratios at 50, 100 and 200 ms ISI in D2 IL-Shell synapses demonstrated that remifentanil treatment significantly increased the paired-pulse ratios at 50 ms ISI. At 100 and 200 ms ISI paired-pulse ratio of males was significantly higher than females. Sal – Saline, Rem – Remifentanil, M – males, F – females; Summary data are presented as Mean ± SEM. \**p* < 0.05 for treatment effects; ^#^*p* < 0.05 for sex effects.

At D2 IL-Shell synapses, there was no significant effect of remifentanil treatment (*F*_(1, 25)_ = 0.406, *p* = 0.530), sex (*F*_(1, 25)_ = 1.239, *p* = 0.276) or interaction between treatment and sex (*F*_(1, 25)_ = 0.005, *p* = 0.947) on A/N ratios (**Fig. 6H, I**). In contrast to D1 IL-Shell synapses, no impact of treatment (*F*_(1, 23)_ = 0.007, *p* = 0.935) or sex (*F*_(1, 23)_ = 1.436, *p* = 0.243), or an interaction of treatment and sex (*F*_(1, 23)_ = 0.428, *p* = 0.519) was observed for AMPAR rectification indices at D2 IL-Shell synapses (**Fig. 6J-L**). Assessment of paired-pulse ratios at D2 IL-Shell synapses showed a significant interaction between treatment and sex across ISIs (*F*_(2, 64)_ = 15.286, *p* < 0.001). Assessment of pairwise comparisons showed that at D2 IL-Shell synapses in males, remifentanil treatment increased paired-pulse ratio at the 50 ms ISI compared to saline (*p* = 0.004). Examination within individual ISIs showed that at 50 ms ISI, a significant interaction between treatment and sex (*F*_(1, 32)_ = 5.922, *p* = 0.021) was observed, with males exhibiting increased paired-pulse ratio following remifentanil treatment vs females. At 100 ms ISI, we found a significant effect of sex (*F*_(1, 32)_ = 4.209, *p* = 0.048) but no significant effect of remifentanil treatment (*F*_(1, 32)_ = 0.429, *p* = 0.517) or an interaction between treatment and sex (*F*_(1, 32)_ = 1.367, *p* = 0.251). At 200 ms ISI, we also found a significant effect of sex (*F*_(1, 32)_ = 7.249, *p* = 0.011) with paired-pulse ratio at D2 IL-Shell synapses of males being significantly greater than females (*p* = 0.011). However, we did not see a significant effect of treatment (*F*_(1, 32)_ = 0.070, *p* = 0.793) or a significant interaction between treatment and sex (*F*_(1, 32)_ = 0.486, *p* = 0.491; **Fig. 6M, N**). Together, these data suggest that decreased presynaptic strength at D2 MSNs is likely due to decreased presynaptic release probability at D2 IL-Shell synapses in males.

## 4 DISCUSSION

Previous research, including our own, has characterized opioid-induced neuroadaptations in NAc glutamate signaling at the level of either specific cell types or subregions (Graziane *et al*., 2016; Hearing *et al*., 2016; Madayag *et al*., 2019). However, examination of sex-, cell type-, and input-specific baseline differences in glutamate signaling as well as whether these variables influence opioid-induced plasticity has yet to be explored. Furthermore, this past work has been confined to non-contingent models of opioid administration. Our study reveals a complex pattern of baseline glutamatergic signaling and synaptic plasticity arising during remifentanil abstinence that is specific to sex, NAc subregion (core vs. shell), and neuronal cell type (D1 vs. D2 MSNs).

### 4.1 Differences in baseline MSN glutamate transmission

We first examined baseline differences in synaptic properties of D1 and D2 MSNs in the NAc core and shell. In the core, D2 MSNs exhibited significantly greater mEPSC amplitude and frequency compared to D1 counterparts in males but not females. While increase in amplitude are likely attributed to enhanced postsynaptic signaling, elevations in frequency may be a result of increased receptor (or synapse) number or increased presynaptic release probability (Kerchner & Nicoll, 2008; Han & Stevens, 2009; Graziane *et al*., 2016; Hearing *et al*., 2016). While previous work has demonstrated higher excitatory drive at D2 MSNs in dorsal striatal regions (Kreitzer & Malenka, 2007), this is the first known evidence that this phenotype is also present in ventral striatal regions. Unlike previous studies in which no baseline differences in glutamate signaling were observed at ventral hippocampus and BLA inputs to the NAc (MacAskill *et al*., 2014), the present study observed cell-type and projection-specific baseline differences in synaptic signaling. We observed lower A/N ratios and AMPAR rectification indices at D2 PL-Core synapses compared to D1 synapses which likely reflect higher levels of CP-AMPAR mediated signaling and greater synaptic strength. We also observed greater levels of release probability at D2 PL-Core vs D1 synapses. The implications of this difference are unclear, it is possible that it represents a baseline “protective” measure that when removed, may lead to maladaptive drug related behavior, as reductions in synaptic strength at D2 NAc core MSNs has been linked to increased habit-like cocaine seeking behavior (Bock *et al*., 2013). Notably, we observed greater levels of release probability at D2 vs D1 PL-Core synapses specifically in males. The lack of differences in cell-type observed in females suggests a sex difference in baseline transmission. In agreement with our findings, past work has shown that MSNs in the NAc core of females have greater spine density (Forlano & Woolley, 2010; Wissman *et al*., 2011) and large spine diameter (Forlano & Woolley, 2010) – findings that align with greater frequency of mEPSCs (Wissman *et al*., 2011). Importantly, the present study is the first to demonstrate that these differences likely exhibit cell-type and input-specificity. Therefore, it is possible that baseline differences in synaptic properties between males and females may underlie sex differences in reward-related disorders or increase vulnerability to transition to use disorders more rapidly (Becker & Hu, 2008; Kniffin & Briand, 2024).

In contrast to the NAc core, baseline differences across cell-types were much more restricted, with higher mEPSC frequency observed at D2 IL-Shell synapses of both males and females. While a trend towards increased A/N ratios at IL-Shell D1 versus D2 synapses, distinctions in presynaptic release were confined to males at these synapses. It is worth noting that the lack of sex differences in the NAc shell agrees with previous pathway non-specific findings that while MSNs in females have larger spines, no differences in MSN intrinsic properties or mEPSCs were present (Willett *et al*., 2016). Regardless, increasing research is demonstrating the influence of sex, and sex hormones on intrinsic and synaptic properties of D1 and D2 MSNs (Cao *et al*., 2018; Proano *et al*., 2018). Thus, future studies controlling for the estrus cycle stage of female mice would help identify the role of sex hormones on the synaptic properties of these neurons.

### 4.2 Cell- and Sex-Specific Neuroadaptations in the NAc Core after remifentanil

The most pronounced differences identified during remifentanil abstinence emerged in the NAc core. In male mice, we observed a pathway non-specific increase in mEPSC amplitude in D1 MSNs – an effect that was not observed in females. However, both sexes exhibited an increase in mEPSC frequency, highlighting an upregulation of excitatory drive that is overlapping and divergent across sex. Alternatively, isolation of changes at PL-Core synapses showed a robust reduction of A/N ratios that aligned with a reduction in the AMPAR rectification indices in remifentanil treated males and females, indicative of increased postsynaptic CP-AMPAR signaling. These data coupled with a decreased paired-pulse ratio, provide convergent evidence for enhanced presynaptic glutamate release and increased postsynaptic AMPA receptor sensitivity that may be more pronounced at PL-Core synapses. Notably, this potentiation of excitatory signaling at D1 NAc MSNs has long been thought to reflect a hallmark of addiction-related neuroplasticity produced by both opioids and psychostimulants (Bock *et al*., 2013; Hikida *et al*., 2016; Hearing *et al*., 2018; Klawonn & Malenka, 2018); however, this is the first indication that this occurs following opioid self-administration. This is particularly pertinent, as pathway non-specific increases at D1 MSNs of NAc core were not previously observed following non-contingent morphine exposure (Hearing *et al*., 2016). While unclear, it is possible that the emergence of plasticity in a more motor-centric region of the NAc reflects the use of an operant- and goal-directed model of drug exposure.

Co-occurring with augmented excitatory drive at D1 MSNs was an observed weakening of glutamate signaling at D2 MSNs. This was prominent when assessing pathway non-specific inputs (e.g., both mEPSC amplitude and frequency). Subsequent examination of PL-Core plasticity showed more nuanced effects, however. Evidenced by an increase in the paired-pulse ratio, presynaptic release was decreased in both males and females. Alternatively, examination of postsynaptic plasticity showed effects selectively in males, with an increase in A/N ratios in remifentanil treated mice. While CP-AMPARs are not traditionally thought to be the predominant sub-unit under baseline conditions, rectification indices were lower and A/N ratios higher at D2 PL-Core synapses compared to D1 synapses in saline treated males. Thus, it is possible that remifentanil promotes a subunit-composition shift by reducing expression of CP-AMPARs and increasing expression of AMPARs that contain the GluA2 subunit. In support, GluA2-containing AMPARs have a roughly linear current-voltage relationship that permits more outward current at positive voltages, which in turn may lead to the observed increase in A/N ratios following remifentanil self-administration. Given that D2 MSNs are part of the "no-go" pathway, which suppresses inappropriate or maladaptive actions (Bock *et al*., 2013; Soares-Cunha *et al*., 2018; Soares-Cunha *et al*., 2020), the simultaneous strengthening of the D1 pathway and weakening of the D2 pathway may reflect a neurobiological bias towards drug-related behavior that arises during abstinence and leaves individuals vulnerable to subsequent relapse.

### 4.2 Cell- and Sex-Specific Neuroadaptations in the NAc Shell after remifentanil

The shell subdivision of the NAc is highly connected with limbic and autonomic brain regions and has been shown to be heavily involved in drug-associated motivation, reward learning, and relapse (Heimer *et al*., 1997; Sesack & Grace, 2010; Bossert *et al*., 2012; Pascoli *et al*., 2014; Hearing *et al*., 2016; Hikida *et al*., 2016). Previous work has shown that at protracted withdrawal timepoints following repeated non-contingent morphine exposure, synaptic strength is increased and decreased at NAc shell D1 and D2 MSNs, respectively (Graziane *et al*., 2016; Hearing *et al*., 2016; Madayag *et al*., 2019). To our knowledge, our study is the first to directly examine the impact of self-administered opioids on synaptic transmission in the NAc shell and how this may differ based on biological sex. Similar to non-contingent exposure, remifentanil increased mEPSC amplitude at D1 MSNs in both sexes, indicative of enhanced postsynaptic AMPAR signaling. Unexpectedly, increases in mEPSC frequency were only observed in males. Additionally, similar to non-contingent morphine, neuroadaptations at D2 MSNs were only observed through decreases in mEPSC frequency – a phenomenon that was previously shown to reflect reduced presynaptic release from pooled afferents (Hearing *et al*., 2016). This divergent regulation of synaptic transmission in D1 vs. D2 MSNs is congruent with their antagonistic roles in addiction behavior and the notion that MSN subpopulations receive differential afferent innervation (Wall *et al*., 2013; MacAskill *et al*., 2014; Boxer *et al*., 2023). The exact mechanism behind cell-type specific plasticity remains unclear; however, past work suggests that increased release at D1 MSNs may reflect a reduction in presynaptic mu opioid receptor inhibition (James *et al*., 2013). Conversely, reduction in release at D2 MSNs reflect alterations in endocannabinoid-mediated LTD (Grueter *et al*., 2010). Regardless, the overlap of these adaptations with non-contingent exposure as well as across sex likely highlight key adaptations associated with opioid motivation and associative learning that modulate behavior.

Previous studies have indirectly highlighted the importance of the IL-Shell pathway in relapse and reinstatement of conditioned place preference (Bossert & Daniel, 2006; Hearing *et al*., 2016; Kruyer *et al*., 2019; Madayag *et al*., 2019; Chioma *et al*., 2021); however, the cell-type and sex-specific nature of adaptations associated with behavior was previously unknown. As seen in the PL-Core circuit, remifentanil produced a decrease in both the A/N ratio and AMPAR rectification at D1 IL-Shell synapses. However, unlike the PL-Core, this effect was only observed in males. Potentiation of this circuit aligns with previous findings following non-contingent morphine and cocaine self-administration in male mice; however, it highlights an intriguing divergence in plasticity across sex. Furthermore, as no effects of remifentanil were observed in the amplitude of mEPSCs in pathway agnostic D1 NAc shell MSNs, it suggests that opioid self-administration may specifically and more prominently impact the IL-Shell network to impact behavior. Indeed, previous work has shown a selective potentiation of IL but not amygdala or ventral hippocampal input at D1 NAc shell MSNs following cocaine self-administration (Pascoli *et al*., 2014) – a phenomenon that is particularly intriguing given that shell MSNs have been shown to receive greater innervation from ventral hippocampus afferents (Britt *et al*., 2012). While the functional implications of these adaptations are unclear, they may underlie increased negative affect and dysphoria during abstinence that motivates continued drug use (Flagel *et al*., 2007; Hikida *et al*., 2016; Madayag *et al*., 2019).

There are several limitations to this study that should be acknowledged. First, food-based lever training and a fading procedure were used to initiate acquisition of self-administration. Although this fading approach was utilized to expedite acquisition of lever pressing and reduce the rate of failed acquisition, it also resulted in an atypical maintenance of lever pressing in saline treated mice (see supplemental materials). We do not believe this reflects a lack of reinforcing properties of remifentanil and sustained responding for Ensure, but rather, a lack of or slowed extinction of responding for Ensure^®^. There are a number of reasons for this: first, past work in mice shows that the average time to extinguish responding for a non-drug reward is ∼10-12 days (Bobadilla *et al*., 2017). Second, previous work from our group using a similar fading approach showed that remifentanil treated mice demonstrated increased motivation compared to saline in a progressive ratio test (Anderson *et al*., 2021). Finally, it has been reported that mice will actively press a lever to produce a light cue in an operant task (Olsen & Winder, 2009). We also do not think that observed differences in physiological properties at baseline, as well as across treatment reflect adaptations associated with extinction in saline mice. This is because in a previous study we did not find differences in physiological properties of mPFC pyramidal neurons between mice that underwent fading followed by saline self-administration compared to behaviorally naïve mice (Anderson *et al*., 2021). Furthermore, baseline mEPSC amplitudes and oEPSCs in the present study are nearly identical to our previous work where mice received non-contingent saline without extinction (Hearing *et al*., 2016). A second caveat of this study is that a non-drug reward was not used for comparison with remifentanil, thus we are unable to determine how selective these adaptations are for opioids. However, we have previously demonstrated that two weeks of Ensure^®^ self-administration did not promote plasticity in mPFC pyramidal neurons (Anderson *et al*., 2021). Finally, our experiments do not address alterations in gonadal hormones. As recent work has shown a role of estrous stage on cue/drug associations (Johnson *et al*., 2019), heroin self-administration (Lacy *et al*., 2016), and remifentanil demand (Lacy *et al*., 2020), a critical step in future studies will be to determine what role, if any, these hormones have on plasticity.

## 5 CONCLUSIONS

Our ability to effectively treat substance use disorders is likely hindered by variability within diagnosed populations. Biological sex is known to dictate drug-related behavior and outcomes, as females exhibit heightened risk for substance use disorders. Data from the present study collectively align with the broader research on sex differences in addiction, which often shows that mechanisms underlying substance abuse may differ between males and females (Becker & Chartoff, 2019; Kokane & Perrotti, 2020; Nicolas *et al*., 2022). While adaptations observed in male and female mice cannot definitively be mapped on to alterations in humans, our findings highlight several potentially key points of overlap produced by opioids as well as other illicit substances that may represent targetable changes across sex and use disorders.

## 6 Conflict of Interest

The authors declare that the research was conducted in the absence of any commercial or financial relationships that could be construed as a potential conflict of interest.

## 7 Author Contributions

M.C.H., A.C.M. and E.M.A. designed experiments; M.C.H., A.C.M., E.M.A., S.D., A.E., and L.F. performed experiments; S.S.K and S.I.A. analyzed data, prepared figures and drafted manuscript; S.S.K, S.I.A. and M.C.H. interpreted results of experiments, edited and revised the manuscript.

## 8 Funding

This study was supported by NIH R00 DA038706-03 and R01 DA055956-01 awarded to M.C.H by the National Institutes of Health’s National Institute for Drug Addiction. The content is solely the responsibility of the authors and does not necessarily represent the official views of the National Institutes of Health’s National Institute for Drug Addiction.

## 9 Acknowledgements

We would like acknowledge Megan Matre for her contributions in conducting behavioral experiments included in this study. We would also like to thank all members of the Hearing laboratory, as well as Dr. Mark Thomas for initial technical insight and equipment support.

## Notes

### Competing Interest Statement

The authors have declared no competing interest.

## REFERENCES

Anderson, E. M., Gomez, D., Caccamise, A., McPhail, D. and Hearing, M. (2019) Chronic unpredictable stress promotes cell-specific plasticity in prefrontal cortex D1 and D2 pyramidal neurons. Neurobiol Stress, 10, 100152. 10.1016/j.ynstr.2019.100152. Available at: https://www.ncbi.nlm.nih.gov/pubmed/30937357.

Anderson, E. M., Engelhardt, A., Demis, S., Porath, E. and Hearing, M. C. (2021) Remifentanil self-administration in mice promotes sex-specific prefrontal cortex dysfunction underlying deficits in cognitive flexibility. Neuropsychopharmacology, 46(10), 1734–1745. 10.1038/s41386-021-01028-z. Available at: https://www.ncbi.nlm.nih.gov/pubmed/34012018.

Becker, J. B. and Hu, M. (2008) Sex differences in drug abuse. Front Neuroendocrinol, 29(1), 36–47. 10.1016/j.yfrne.2007.07.003. Available at: https://www.ncbi.nlm.nih.gov/pubmed/17904621.

Becker, J. B. and Chartoff, E. (2019) Sex differences in neural mechanisms mediating reward and addiction. Neuropsychopharmacology, 44(1), 166–183. 10.1038/s41386-018-0125-6. Available at: https://www.ncbi.nlm.nih.gov/pubmed/29946108.

Berendse, G.-d. G., Groenewegen (1992) Topographical Organization and Relationship with Ventral Striatal Compartments of Prefrontal Corticostriatal Projections in the Rat. Journal of Comparative Neurology.

Bobadilla, A. C., Garcia-Keller, C., Heinsbroek, J. A., Scofield, M. D., Chareunsouk, V., Monforton, C. and Kalivas, P. W. (2017) Accumbens Mechanisms for Cued Sucrose Seeking. Neuropsychopharmacology, 42(12), 2377–2386. 10.1038/npp.2017.153. Available at: https://www.ncbi.nlm.nih.gov/pubmed/28726801.

Bock, R., Shin, J. H., Kaplan, A. R., Dobi, A., Markey, E., Kramer, P. F., Gremel, C. M., Christensen, C. H., Adrover, M. F. and Alvarez, V. A. (2013) Strengthening the accumbal indirect pathway promotes resilience to compulsive cocaine use. Nat Neurosci, 16(5), 632–8. 10.1038/nn.3369. Available at: https://www.ncbi.nlm.nih.gov/pubmed/23542690.

Bossert, J. and Daniel, C. (2006) trans-cis Photoisomerization of the styrylpyridine Ligand in [Re(CO)3(2,2’-bipyridine)(t-4-styrylpyridine)]+: role of the metal-to-ligand charge-transfer excited states. Chemistry, 12(18), 4835–43. 10.1002/chem.200501082. Available at: https://www.ncbi.nlm.nih.gov/pubmed/16642521.

Bossert, J. M., Stern, A. L., Theberge, F. R., Marchant, N. J., Wang, H. L., Morales, M. and Shaham, Y. (2012) Role of projections from ventral medial prefrontal cortex to nucleus accumbens shell in context-induced reinstatement of heroin seeking. J Neurosci, 32(14), 4982–91. 10.1523/JNEUROSCI.0005-12.2012. Available at: https://www.ncbi.nlm.nih.gov/pubmed/22492053.

Boxer, E. E., Kim, J., Dunn, B. and Aoto, J. (2023) Ventral Subiculum Inputs to Nucleus Accumbens Medial Shell Preferentially Innervate D2R Medium Spiny Neurons and Contain Calcium Permeable AMPARs. J Neurosci, 43(7), 1166–1177. 10.1523/JNEUROSCI.1907-22.2022. Available at: https://www.ncbi.nlm.nih.gov/pubmed/36609456.

Britt, J. P., Benaliouad, F., McDevitt, R. A., Stuber, G. D., Wise, R. A. and Bonci, A. (2012) Synaptic and behavioral profile of multiple glutamatergic inputs to the nucleus accumbens. Neuron, 76(4), 790–803. 10.1016/j.neuron.2012.09.040. Available at: https://www.ncbi.nlm.nih.gov/pubmed/23177963.

Brog, J. S., Salyapongse, A., Deutch, A. Y. and Zahm, D. S. (1993) The patterns of afferent innervation of the core and shell in the "accumbens" part of the rat ventral striatum: immunohistochemical detection of retrogradely transported fluoro-gold. J Comp Neurol, 338(2), 255–78. 10.1002/cne.903380209. Available at: https://www.ncbi.nlm.nih.gov/pubmed/8308171.

Calipari, E. S., Bagot, R. C., Purushothaman, I., Davidson, T. J., Yorgason, J. T., Pena, C. J., Walker, D. M., Pirpinias, S. T., Guise, K. G., Ramakrishnan, C., Deisseroth, K. and Nestler, E. J. (2016) In vivo imaging identifies temporal signature of D1 and D2 medium spiny neurons in cocaine reward. Proc Natl Acad Sci U S A, 113(10), 2726–31. 10.1073/pnas.1521238113. Available at: https://www.ncbi.nlm.nih.gov/pubmed/26831103.

Cao, J., Dorris, D. M. and Meitzen, J. (2018) Electrophysiological properties of medium spiny neurons in the nucleus accumbens core of prepubertal male and female Drd1a-tdTomato line 6 BAC transgenic mice. J Neurophysiol, 120(4), 1712–1727. 10.1152/jn.00257.2018. Available at: https://www.ncbi.nlm.nih.gov/pubmed/29975170.

Chioma, V. C., Kruyer, A., Bobadilla, A. C., Angelis, A., Ellison, Z., Hodebourg, R., Scofield, M. D. and Kalivas, P. W. (2021) Heroin Seeking and Extinction From Seeking Activate Matrix Metalloproteinases at Synapses on Distinct Subpopulations of Accumbens Cells. Biol Psychiatry, 89(10), 947–958. 10.1016/j.biopsych.2020.12.004. Available at: https://www.ncbi.nlm.nih.gov/pubmed/33579535.

Dittman, J. S., Kreitzer, A. C. and Regehr, W. G. (2000) Interplay between facilitation, depression, and residual calcium at three presynaptic terminals. J Neurosci, 20(4), 1374–85. 10.1523/JNEUROSCI.20-04-01374.2000. Available at: https://www.ncbi.nlm.nih.gov/pubmed/10662828.

Everitt, B. J. and Robbins, T. W. (2005) Neural systems of reinforcement for drug addiction: from actions to habits to compulsion. Nat Neurosci, 8(11), 1481–9. 10.1038/nn1579. Available at: https://www.ncbi.nlm.nih.gov/pubmed/16251991.

Flagel, S. B., Watson, S. J., Robinson, T. E. and Akil, H. (2007) Individual differences in the propensity to approach signals vs goals promote different adaptations in the dopamine system of rats. Psychopharmacology (Berl), 191(3), 599–607. 10.1007/s00213-006-0535-8. Available at: https://www.ncbi.nlm.nih.gov/pubmed/16972103.

Forlano, P. M. and Woolley, C. S. (2010) Quantitative analysis of pre- and postsynaptic sex differences in the nucleus accumbens. J Comp Neurol, 518(8), 1330–48. 10.1002/cne.22279. Available at: https://www.ncbi.nlm.nih.gov/pubmed/20151363.

Gerfen, C. R. and Surmeier, D. J. (2011) Modulation of striatal projection systems by dopamine. Annu Rev Neurosci, 34, 441–66. 10.1146/annurev-neuro-061010-113641. Available at: https://www.ncbi.nlm.nih.gov/pubmed/21469956.

Glass, M. J., Lane, D. A., Colago, E. E., Chan, J., Schlussman, S. D., Zhou, Y., Kreek, M. J. and Pickel, V. M. (2008) Chronic administration of morphine is associated with a decrease in surface AMPA GluR1 receptor subunit in dopamine D1 receptor expressing neurons in the shell and non-D1 receptor expressing neurons in the core of the rat nucleus accumbens. Exp Neurol, 210(2), 750–61. 10.1016/j.expneurol.2008.01.012. Available at: https://www.ncbi.nlm.nih.gov/pubmed/18294632.

Graziane, N. M., Sun, S., Wright, W. J., Jang, D., Liu, Z., Huang, Y. H., Nestler, E. J., Wang, Y. T., Schluter, O. M. and Dong, Y. (2016) Opposing mechanisms mediate morphine- and cocaine-induced generation of silent synapses. Nat Neurosci, 19(7), 915–25. 10.1038/nn.4313. Available at: https://www.ncbi.nlm.nih.gov/pubmed/27239940.

Grueter, B. A., Brasnjo, G. and Malenka, R. C. (2010) Postsynaptic TRPV1 triggers cell type-specific long-term depression in the nucleus accumbens. Nat Neurosci, 13(12), 1519–25. 10.1038/nn.2685. Available at: https://www.ncbi.nlm.nih.gov/pubmed/21076424.

Han, E. B. and Stevens, C. F. (2009) Development regulates a switch between post- and presynaptic strengthening in response to activity deprivation. Proc Natl Acad Sci U S A, 106(26), 10817–22. 10.1073/pnas.0903603106. Available at: https://www.ncbi.nlm.nih.gov/pubmed/19509338.

Hearing, M., Graziane, N., Dong, Y. and Thomas, M. J. (2018) Opioid and Psychostimulant Plasticity: Targeting Overlap in Nucleus Accumbens Glutamate Signaling. Trends Pharmacol Sci, 39(3), 276–294. 10.1016/j.tips.2017.12.004. Available at: https://www.ncbi.nlm.nih.gov/pubmed/29338873.

Hearing, M. (2019) Prefrontal-accumbens opioid plasticity: Implications for relapse and dependence. Pharmacol Res, 139, 158–165. 10.1016/j.phrs.2018.11.012. Available at: https://www.ncbi.nlm.nih.gov/pubmed/30465850.

Hearing, M. C., Jedynak, J., Ebner, S. R., Ingebretson, A., Asp, A. J., Fischer, R. A., Schmidt, C., Larson, E. B. and Thomas, M. J. (2016) Reversal of morphine-induced cell-type-specific synaptic plasticity in the nucleus accumbens shell blocks reinstatement. Proc Natl Acad Sci U S A, 113(3), 757–62. 10.1073/pnas.1519248113. Available at: https://www.ncbi.nlm.nih.gov/pubmed/26739562.

Heidbreder, C. A. and Groenewegen, H. J. (2003) The medial prefrontal cortex in the rat: evidence for a dorso-ventral distinction based upon functional and anatomical characteristics. Neurosci Biobehav Rev, 27(6), 555–79. 10.1016/j.neubiorev.2003.09.003. Available at: https://www.ncbi.nlm.nih.gov/pubmed/14599436.

Heimer, L., Alheid, G. F., de Olmos, J. S., Groenewegen, H. J., Haber, S. N., Harlan, R. E. and Zahm, D. S. (1997) The accumbens: beyond the core-shell dichotomy. J Neuropsychiatry Clin Neurosci, 9(3), 354–81. 10.1176/jnp.9.3.354. Available at: https://www.ncbi.nlm.nih.gov/pubmed/9276840.

Heimer, L. (2003) The legacy of the silver methods and the new anatomy of the basal forebrain: implications for neuropsychiatry and drug abuse. Scand J Psychol, 44(3), 189–201. 10.1111/1467-9450.00336. Available at: https://www.ncbi.nlm.nih.gov/pubmed/12914582.

Hikida, T., Morita, M. and Macpherson, T. (2016) Neural mechanisms of the nucleus accumbens circuit in reward and aversive learning. Neurosci Res, 108, 1–5. 10.1016/j.neures.2016.01.004. Available at: https://www.ncbi.nlm.nih.gov/pubmed/26827817.

Hoover, W. B. and Vertes, R. P. (2007) Anatomical analysis of afferent projections to the medial prefrontal cortex in the rat. Brain Struct Funct, 212(2), 149–79. 10.1007/s00429-007-0150-4. Available at: https://www.ncbi.nlm.nih.gov/pubmed/17717690.

Hutcheson, D. M., Parkinson, J. A., Robbins, T. W. and Everitt, B. J. (2001) The effects of nucleus accumbens core and shell lesions on intravenous heroin self-administration and the acquisition of drug-seeking behaviour under a second-order schedule of heroin reinforcement. Psychopharmacology (Berl), 153(4), 464–72. 10.1007/s002130000635. Available at: https://www.ncbi.nlm.nih.gov/pubmed/11243494.

James, A. S., Chen, J. Y., Cepeda, C., Mittal, N., Jentsch, J. D., Levine, M. S., Evans, C. J. and Walwyn, W. (2013) Opioid self-administration results in cell-type specific adaptations of striatal medium spiny neurons. Behav Brain Res, 256, 279–83. 10.1016/j.bbr.2013.08.009. Available at: https://www.ncbi.nlm.nih.gov/pubmed/23968589.

Johnson, A. R., Thibeault, K. C., Lopez, A. J., Peck, E. G., Sands, L. P., Sanders, C. M., Kutlu, M. G. and Calipari, E. S. (2019) Cues play a critical role in estrous cycle-dependent enhancement of cocaine reinforcement. Neuropsychopharmacology, 44(7), 1189–1197. 10.1038/s41386-019-0320-0. Available at: https://www.ncbi.nlm.nih.gov/pubmed/30728447.

Kalivas, P. W. (2009) The glutamate homeostasis hypothesis of addiction. Nat Rev Neurosci, 10(8), 561–72. 10.1038/nrn2515. Available at: https://www.ncbi.nlm.nih.gov/pubmed/19571793.

Kerchner, G. A. and Nicoll, R. A. (2008) Silent synapses and the emergence of a postsynaptic mechanism for LTP. Nat Rev Neurosci, 9(11), 813–25. 10.1038/nrn2501. Available at: https://www.ncbi.nlm.nih.gov/pubmed/18854855.

Klawonn, A. M. and Malenka, R. C. (2018) Nucleus Accumbens Modulation in Reward and Aversion. Cold Spring Harb Symp Quant Biol, 83, 119–129. 10.1101/sqb.2018.83.037457. Available at: https://www.ncbi.nlm.nih.gov/pubmed/30674650.

Kniffin, A. R. and Briand, L. A. (2024) Sex differences in glutamate transmission and plasticity in reward related regions. Front Behav Neurosci, 18, 1455478.10.3389/fnbeh.2024.1455478. Available at: https://www.ncbi.nlm.nih.gov/pubmed/39359325.

Kokane, S. S. and Perrotti, L. I. (2020) Sex Differences and the Role of Estradiol in Mesolimbic Reward Circuits and Vulnerability to Cocaine and Opiate Addiction. Front Behav Neurosci, 14, 74. 10.3389/fnbeh.2020.00074. Available at: https://www.ncbi.nlm.nih.gov/pubmed/32508605.

Kokane, S. S., Cole, R. D., Bordieanu, B., Ray, C. M., Haque, I. A., Otis, J. M. and McGinty, J. F. (2023) Increased Excitability and Synaptic Plasticity of Drd1- and Drd2-Expressing Prelimbic Neurons Projecting to Nucleus Accumbens after Heroin Abstinence Are Reversed by Cue-Induced Relapse and Protein Kinase A Inhibition. J Neurosci, 43(22), 4019–4032. 10.1523/JNEUROSCI.0108-23.2023. Available at: https://www.ncbi.nlm.nih.gov/pubmed/37094933.

Koob, G. F., Ahmed, S. H., Boutrel, B., Chen, S. A., Kenny, P. J., Markou, A., O’Dell, L. E., Parsons, L. H. and Sanna, P. P. (2004) Neurobiological mechanisms in the transition from drug use to drug dependence. Neurosci Biobehav Rev, 27(8), 739–49. 10.1016/j.neubiorev.2003.11.007. Available at: https://www.ncbi.nlm.nih.gov/pubmed/15019424.

Koob, G. F. and Volkow, N. D. (2010) Neurocircuitry of addiction. Neuropsychopharmacology, 35(1), 217–38. 10.1038/npp.2009.110. Available at: https://www.ncbi.nlm.nih.gov/pubmed/19710631.

Koob, G. F. and Volkow, N. D. (2016) Neurobiology of addiction: a neurocircuitry analysis. Lancet Psychiatry, 3(8), 760–773. 10.1016/S2215-0366(16)00104-8. Available at: https://www.ncbi.nlm.nih.gov/pubmed/27475769.

Kravitz, A. V., Tye, L. D. and Kreitzer, A. C. (2012) Distinct roles for direct and indirect pathway striatal neurons in reinforcement. Nat Neurosci, 15(6), 816–8. 10.1038/nn.3100. Available at: https://www.ncbi.nlm.nih.gov/pubmed/22544310.

Kreitzer, A. C. and Malenka, R. C. (2007) Endocannabinoid-mediated rescue of striatal LTD and motor deficits in Parkinson’s disease models. Nature, 445(7128), 643–7. 10.1038/nature05506. Available at: https://www.ncbi.nlm.nih.gov/pubmed/17287809.

Kruyer, A., Scofield, M. D., Wood, D., Reissner, K. J. and Kalivas, P. W. (2019) Heroin Cue-Evoked Astrocytic Structural Plasticity at Nucleus Accumbens Synapses Inhibits Heroin Seeking. Biol Psychiatry, 86(11), 811–819. 10.1016/j.biopsych.2019.06.026. Available at: https://www.ncbi.nlm.nih.gov/pubmed/31495448.

Lacy, R. T., Strickland, J. C., Feinstein, M. A., Robinson, A. M. and Smith, M. A. (2016) The effects of sex, estrous cycle, and social contact on cocaine and heroin self-administration in rats. Psychopharmacology (Berl), 233(17), 3201–10. 10.1007/s00213-016-4368-9. Available at: https://www.ncbi.nlm.nih.gov/pubmed/27370020.

Lacy, R. T., Austin, B. P. and Strickland, J. C. (2020) The influence of sex and estrous cyclicity on cocaine and remifentanil demand in rats. Addict Biol, 25(1), e12716. 10.1111/adb.12716. Available at: https://www.ncbi.nlm.nih.gov/pubmed/30779409.

LaLumiere, R. T. and Kalivas, P. W. (2008) Glutamate release in the nucleus accumbens core is necessary for heroin seeking. J Neurosci, 28(12), 3170–7. 10.1523/JNEUROSCI.5129-07.2008. Available at: https://www.ncbi.nlm.nih.gov/pubmed/18354020.

Lefevre, E. M., Gauthier, E. A., Bystrom, L. L., Scheunemann, J. and Rothwell, P. E. (2023) Differential Patterns of Synaptic Plasticity in the Nucleus Accumbens Caused by Continuous and Interrupted Morphine Exposure. J Neurosci, 43(2), 308–318. 10.1523/JNEUROSCI.0595-22.2022. Available at: https://www.ncbi.nlm.nih.gov/pubmed/36396404.

Lobo, M. K., Covington, H. E., 3rd, Chaudhury, D., Friedman, A. K., Sun, H., Damez-Werno, D., Dietz, D. M., Zaman, S., Koo, J. W., Kennedy, P. J., Mouzon, E., Mogri, M., Neve, R. L., Deisseroth, K., Han, M. H. and Nestler, E. J. (2010) Cell type-specific loss of BDNF signaling mimics optogenetic control of cocaine reward. Science, 330(6002), 385–90. 10.1126/science.1188472. Available at: https://www.ncbi.nlm.nih.gov/pubmed/20947769.

MacAskill, A. F., Cassel, J. M. and Carter, A. G. (2014) Cocaine exposure reorganizes cell type- and input-specific connectivity in the nucleus accumbens. Nat Neurosci, 17(9), 1198–207. 10.1038/nn.3783. Available at: https://www.ncbi.nlm.nih.gov/pubmed/25108911.

Madayag, A. C., Gomez, D., Anderson, E. M., Ingebretson, A. E., Thomas, M. J. and Hearing, M. C. (2019) Cell-type and region-specific nucleus accumbens AMPAR plasticity associated with morphine reward, reinstatement, and spontaneous withdrawal. Brain Struct Funct, 224(7), 2311–2324. 10.1007/s00429-019-01903-y. Available at: https://www.ncbi.nlm.nih.gov/pubmed/31201496.

Marchant, N. J., Kaganovsky, K., Shaham, Y. and Bossert, J. M. (2016) Role of corticostriatal circuits in context-induced reinstatement of drug seeking. Brain Res, 1628(Pt A), 219–32. 10.1016/j.brainres.2014.09.004. Available at: https://www.ncbi.nlm.nih.gov/pubmed/25199590.

McFarland, K., Lapish, C. C. and Kalivas, P. W. (2003) Prefrontal glutamate release into the core of the nucleus accumbens mediates cocaine-induced reinstatement of drug-seeking behavior. J Neurosci, 23(8), 3531–7. 10.1523/JNEUROSCI.23-08-03531.2003. Available at: https://www.ncbi.nlm.nih.gov/pubmed/12716962.

Nelson, A. B., Hang, G. B., Grueter, B. A., Pascoli, V., Luscher, C., Malenka, R. C. and Kreitzer, A. C. (2012) A comparison of striatal-dependent behaviors in wild-type and hemizygous Drd1a and Drd2 BAC transgenic mice. J Neurosci, 32(27), 9119–23. 10.1523/JNEUROSCI.0224-12.2012. Available at: https://www.ncbi.nlm.nih.gov/pubmed/22764221.

Nicolas, C., Zlebnik, N. E., Farokhnia, M., Leggio, L., Ikemoto, S. and Shaham, Y. (2022) Sex Differences in Opioid and Psychostimulant Craving and Relapse: A Critical Review. Pharmacol Rev, 74(1), 119–140. 10.1124/pharmrev.121.000367. Available at: https://www.ncbi.nlm.nih.gov/pubmed/34987089.

Olsen, C. M. and Winder, D. G. (2009) Operant sensation seeking engages similar neural substrates to operant drug seeking in C57 mice. Neuropsychopharmacology, 34(7), 1685–94. 10.1038/npp.2008.226. Available at: https://www.ncbi.nlm.nih.gov/pubmed/19145223.

Pascoli, V., Turiault, M. and Luscher, C. (2011) Reversal of cocaine-evoked synaptic potentiation resets drug-induced adaptive behaviour. Nature, 481(7379), 71–5. 10.1038/nature10709. Available at: https://www.ncbi.nlm.nih.gov/pubmed/22158102.

Pascoli, V., Terrier, J., Espallergues, J., Valjent, E., O’Connor, E. C. and Luscher, C. (2014) Contrasting forms of cocaine-evoked plasticity control components of relapse. Nature, 509(7501), 459–64. 10.1038/nature13257. Available at: https://www.ncbi.nlm.nih.gov/pubmed/24848058.

Pecina, S. and Berridge, K. C. (2005) Hedonic hot spot in nucleus accumbens shell: where do mu-opioids cause increased hedonic impact of sweetness? J Neurosci, 25(50), 11777–10.1523/JNEUROSCI.2329-05.2005. Available at: https://www.ncbi.nlm.nih.gov/pubmed/16354936.

Peters, J., Pattij, T. and De Vries, T. J. (2013) Targeting cocaine versus heroin memories: divergent roles within ventromedial prefrontal cortex. Trends Pharmacol Sci, 34(12), 689–95. 10.1016/j.tips.2013.10.004. Available at: https://www.ncbi.nlm.nih.gov/pubmed/24182624.

Pontieri, F. E., Tanda, G. and Di Chiara, G. (1995) Intravenous cocaine, morphine, and amphetamine preferentially increase extracellular dopamine in the "shell" as compared with the "core" of the rat nucleus accumbens. Proc Natl Acad Sci U S A, 92(26), 12304–8. 10.1073/pnas.92.26.12304. Available at: https://www.ncbi.nlm.nih.gov/pubmed/8618890.

Proano, S. B., Morris, H. J., Kunz, L. M., Dorris, D. M. and Meitzen, J. (2018) Estrous cycle-induced sex differences in medium spiny neuron excitatory synaptic transmission and intrinsic excitability in adult rat nucleus accumbens core. J Neurophysiol, 120(3), 1356– 1373. 10.1152/jn.00263.2018. Available at: https://www.ncbi.nlm.nih.gov/pubmed/29947588.

Rogers, J. L., Ghee, S. and See, R. E. (2008) The neural circuitry underlying reinstatement of heroin-seeking behavior in an animal model of relapse. Neuroscience, 151(2), 579–88. 10.1016/j.neuroscience.2007.10.012. Available at: https://www.ncbi.nlm.nih.gov/pubmed/18061358.

Scofield, M. D., Heinsbroek, J. A., Gipson, C. D., Kupchik, Y. M., Spencer, S., Smith, A. C., Roberts-Wolfe, D. and Kalivas, P. W. (2016) The Nucleus Accumbens: Mechanisms of Addiction across Drug Classes Reflect the Importance of Glutamate Homeostasis. Pharmacol Rev, 68(3), 816–71. 10.1124/pr.116.012484. Available at: https://www.ncbi.nlm.nih.gov/pubmed/27363441.

Sesack, S. R., Deutch, A. Y., Roth, R. H. and Bunney, B. S. (1989) Topographical organization of the efferent projections of the medial prefrontal cortex in the rat: an anterograde tract-tracing study with Phaseolus vulgaris leucoagglutinin. J Comp Neurol, 290(2), 213–42. 10.1002/cne.902900205. Available at: https://www.ncbi.nlm.nih.gov/pubmed/2592611.

Sesack, S. R. and Grace, A. A. (2010) Cortico-Basal Ganglia reward network: microcircuitry. Neuropsychopharmacology, 35(1), 27–47. 10.1038/npp.2009.93. Available at: https://www.ncbi.nlm.nih.gov/pubmed/19675534.

Shen, H., Moussawi, K., Zhou, W., Toda, S. and Kalivas, P. W. (2011) Heroin relapse requires long-term potentiation-like plasticity mediated by NMDA2b-containing receptors. Proc Natl Acad Sci U S A, 108(48), 19407–12. 10.1073/pnas.1112052108. Available at: https://www.ncbi.nlm.nih.gov/pubmed/22084102.

Soares-Cunha, C., Coimbra, B., Domingues, A. V., Vasconcelos, N., Sousa, N. and Rodrigues, A. J. (2018) Nucleus Accumbens Microcircuit Underlying D2-MSN-Driven Increase in Motivation. eNeuro, 5(2). 10.1523/ENEURO.0386-18.2018. Available at: https://www.ncbi.nlm.nih.gov/pubmed/29780881.

Soares-Cunha, C., de Vasconcelos, N. A. P., Coimbra, B., Domingues, A. V., Silva, J. M., Loureiro-Campos, E., Gaspar, R., Sotiropoulos, I., Sousa, N. and Rodrigues, A. J. (2020) Nucleus accumbens medium spiny neurons subtypes signal both reward and aversion. Mol Psychiatry, 25(12), 3241–3255. 10.1038/s41380-019-0484-3. Available at: https://www.ncbi.nlm.nih.gov/pubmed/31462765.

Van den Oever, M. C., Spijker, S., Smit, A. B. and De Vries, T. J. (2010) Prefrontal cortex plasticity mechanisms in drug seeking and relapse. Neurosci Biobehav Rev, 35(2), 276–84. 10.1016/j.neubiorev.2009.11.016. Available at: https://www.ncbi.nlm.nih.gov/pubmed/19932711.

van Zessen, R., Li, Y., Marion-Poll, L., Hulo, N., Flakowski, J. and Luscher, C. (2021) Dynamic dichotomy of accumbal population activity underlies cocaine sensitization. Elife, 10. 10.7554/eLife.66048. Available at: https://www.ncbi.nlm.nih.gov/pubmed/34608866.

Vertes, R. P. (2004) Differential projections of the infralimbic and prelimbic cortex in the rat. Synapse, 51(1), 32–58. 10.1002/syn.10279. Available at: https://www.ncbi.nlm.nih.gov/pubmed/14579424.

Wall, N. R., De La Parra, M., Callaway, E. M. and Kreitzer, A. C. (2013) Differential innervation of direct- and indirect-pathway striatal projection neurons. Neuron, 79(2), 347–60. 10.1016/j.neuron.2013.05.014. Available at: https://www.ncbi.nlm.nih.gov/pubmed/23810541.

Willett, J. A., Will, T., Hauser, C. A., Dorris, D. M., Cao, J. and Meitzen, J. (2016) No Evidence for Sex Differences in the Electrophysiological Properties and Excitatory Synaptic Input onto Nucleus Accumbens Shell Medium Spiny Neurons. eNeuro, 3(1). 10.1523/ENEURO.0147-15.2016. Available at: https://www.ncbi.nlm.nih.gov/pubmed/27022621.

Wissman, A. M., McCollum, A. F., Huang, G. Z., Nikrodhanond, A. A. and Woolley, C. S. (2011) Sex differences and effects of cocaine on excitatory synapses in the nucleus accumbens. Neuropharmacology, 61(1-2), 217–27. 10.1016/j.neuropharm.2011.04.002. Available at: https://www.ncbi.nlm.nih.gov/pubmed/21510962.

Zahm, D. S. and Brog, J. S. (1992) On the significance of subterritories in the "accumbens" part of the rat ventral striatum. Neuroscience, 50(4), 751–67. 10.1016/0306-4522(92)90202-d. Available at: https://www.ncbi.nlm.nih.gov/pubmed/1448200.

Zhu, Y., Wienecke, C. F., Nachtrab, G. and Chen, X. (2016) A thalamic input to the nucleus accumbens mediates opiate dependence. Nature, 530(7589), 219–22. 10.1038/nature16954. Available at: https://www.ncbi.nlm.nih.gov/pubmed/26840481.

