## Supplementary Materials for "Remifentanil self-administration promotes circuit- and sex-specific adaptations within the prefrontal-accumbens pathways"

**SUPPLEMENTARY FIGURES & LEGENDS**

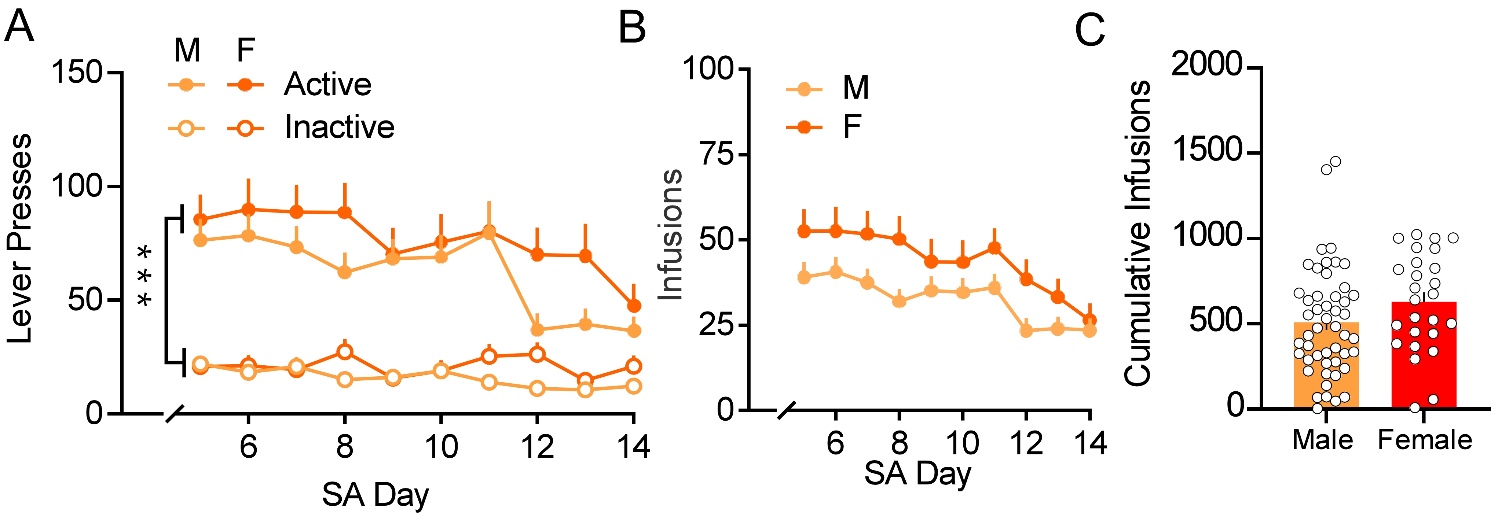

**Supplemental 1. Saline self-administration in males and females.**

**A**. Summary graph of lever pressing behavior over 5-14 days of saline SA. Males (light orange circles) and females (dark orange circles) had significantly greater number of active lever presses (solid circles) vs inactive lever presses (bordered circles). **B**. Summary graph of average saline infusions. Males (light orange squares) and females (dark orange squares) did not differ in daily number of saline infusions. **C**. Cumulative infusions of saline in males (light orange) and females (dark orange). Summary data are presented as Mean ± SEM. ****p <* .001 (Active vs Inactive)*.*
